# Structure-based prediction of Ras-effector binding affinities and design of ‘branchegetic’ interface mutations

**DOI:** 10.1101/2022.09.04.506480

**Authors:** Philipp Junk, Christina Kiel

**Affiliations:** Department of Molecular Medicine, University of Pavia, 27100 Pavia, Italy; Systems Biology Ireland, School of Medicine, University College Dublin, Dublin 4, Ireland; UCD Charles Institute of Dermatology, School of Medicine, University College Dublin, Dublin 4, Ireland

**Keywords:** AlphaFold, branch pruning, edgetics, enedgetics, effectors, FoldX, homology modelling, hub, networks, Ras

## Abstract

Ras is a central cellular hub protein controlling multiple cell fates. How Ras interacts with a variety of potential effector proteins is relatively unexplored, with only some key effectors characterized in great detail. Here, we have used homology modelling based on X-ray and AlphaFold2 templates to build structural models for 54 Ras-effectors complexes. These models were used to estimate binding affinities using a supervised learning regressor. Furthermore, we systematically introduced Ras ‘branch-pruning’ (or branchegetic) mutations to identify 200 interface mutations that affect the binding energy with at least one of the model structures. The impacts of these branchegetic mutants were integrated into a mathematical model to assess the potential for rewiring interactions at the Ras hub on a systems level. These findings have provided a quantitative understanding of Ras-effector interfaces and their impact on systems properties of a key cellular hub.

## Introduction

Ras is a key cellular signaling hub and oncogene ^1^. The first correct Ras structure was famously described in 1989 ^2 3^ and consists of the G domain super-fold (six β-strands and five α-helices; ^1^). It’s main structural and functional characteristic is the nucleotide binding site (the typical α,β-fold of nucleotide binding proteins), which can bind GTP and hydrolyze it to GDP. Then, GDP gets released and a new nucleotide (favorably GTP since it is higher abundant in cells) gets bound. Depending on which nucleotide is bound, the functional state of RAS is different. GDP bound Ras assumes a so-called inactive conformation, whereas upon the binding of GTP, the conformation of Ras is called active. The difference between the active and the inactive conformation is that, in the active conformation, two loop regions called switch 1 and switch 2 are interacting with the GTP and thereby tightly bound to it ^1^ (Figure 1A). This rearranges the interface of Ras in a specific way so that effector proteins with a specific structural motive can interact with Ras. The structural motive required for the interaction with Ras has a ubiquitin domain super-fold ^4^. There exist three families of these domains, called Ras binding domain (RBD), Ras association domain (RA domain) and PI3K-Ras binding domain (PI3K RBD) with only minor sequence and structural differences between them. In the following, for the sake of simplicity, all of these will be called RBDs.

**Figure 1.**
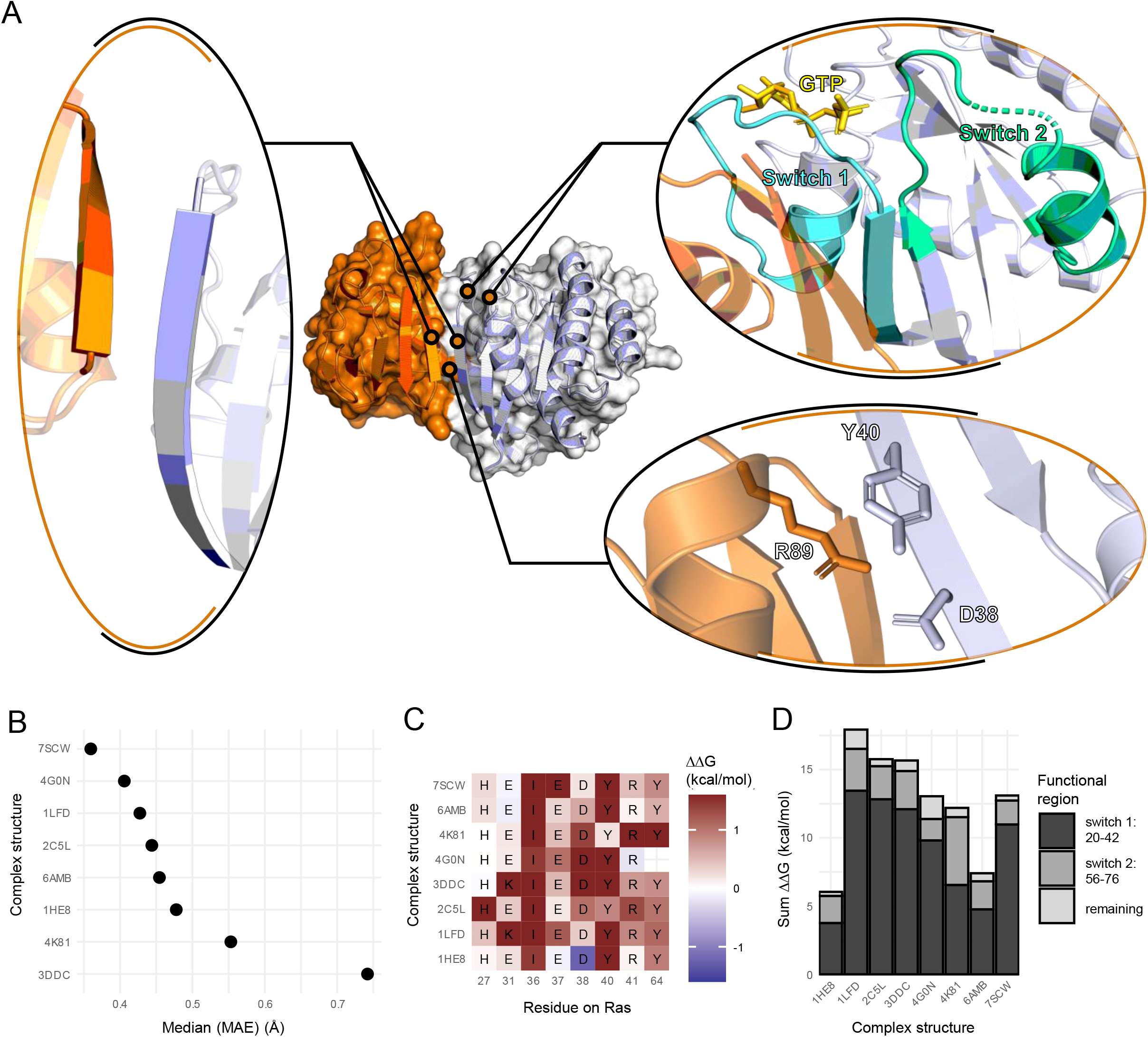
Noticeable features of the Ras-RBD interfaces. (A) Overview of interface features for the Ras-RAF1 interface. Highlighted are the intramolecular β-sheet alignment, the assembly of the Ras interface by the switch regions and energetic hotspots in the interface. (B) β-sheet alignment for experimentally determined complexes (MAE). A lower value indicates a more similar alignment of the intramolecular β-sheets compared to other crystal structures. (C) FoldX alanine scan hotspots on RAS for experimentally determined complex structures. The color scale is confined to the limits [-1.6, 1.6]. Positive ΔΔG values indicate a loss in binding energy upon alanine mutation, which reflects that the respective amino acid contributes to binding. (D) Energetic contributions of functional regions based on alanine scan analysis.

Ras·GTP can interact and activate many downstream effectors, some of which are better studied than others ^5^. Well-studied effectors include PI3-kinases, RalGDS, and Raf kinases, which control important cellular processes such as survival, polarization, adhesion, migration, and proliferation. In our previous work, we characterized a set of 56 RBD-containing proteins as potential Ras effectors, which converge into 12 classes that are linked to different downstream cellular processes and phenotypes ^6 7 8 9^.

Ras-effector interactions are interesting from a systems biology and network point of view. Effectors use a mutually exclusive binding site on Ras, hence competition for binding can occur under certain conditions ^10^. Also, the experimental and predicted affinities between RBDs and Ras·GTP vary, ranging from nano- to micromolar K_d_ values ^8^. Previously we analyzed how the amount of effectors in complex with Ras varies in different cell types ^6 7^. We find that only 9 effectors are predicted to bind in significant amount to Ras·GTP using the RBD alone ^7^. However, 31 effectors are predicted to form significant complexes with Ras if they are additionally recruited to the plasma membrane via other domains present in effectors (piggyback mechanism, ^11^). Hence, even weak binding affinities on the level of RBD-RAS binding can turn into significant complex formation if effectors are recruited (e.g. in a context specific manner). Thus, for systems analyses and computational models, we need good estimations for all binding affinities between RBDs and Ras.

Structural analysis can lead to a deeper understanding of protein-protein interaction (PPI). Previous we used homology modelling, based on experimentally determined complex structures, to predict binding between effectors and Ras·GTP ^12 13^. While this has provided insights into binding propensities, a limitation of this work was that no quantitative binding affinities were predicted, but qualitative binding by classifying effectors into categories of ‘binding’, ‘non-binding’, and ‘twilight’. It is now timely to revisit homology modelling of Ras-RBD complexes, as new template structures are available by (i) X-ray crystallography (8 of 56 effectors are crystallized in complex with RAS) and (ii) AlphaFold ^14 15^.

Ras is frequently mutated in different cancers, where aberrant Ras signaling plays a role in cancer initiation, progression, and metastasis via alterations in metabolism, proliferation, and survival ^16^. As Ras cannot be directly targeted (or only certain Ras mutations) ^17^, much hope rests on network-centric approaches ^18^ that involve Ras-effector interactions or downstream pathways ^9^. Therefore, tinkering with Ras-effector binding is an attractive alternative strategy for finding suitable targets for therapeutic interventions. Previously we developed a ‘branch-pruning’ (or “branchegetic”, in analogy to “edgetics” ^19^ and “enedgetics” ^20^) strategy for Ras effector interactions, where mutations are introduced into Ras that differentially impact binding to effectors ^21^. For example, introducing a mutation can result in a steric clash in the interface formed with one effector, but not with another; hence the interaction with one effector is broken while intact for another.

In this work, we first generate homology models of all Ras-effector (RBD) interactions and predict affinities of the RBDs in complex with Ras·GTP. We then use the generated model structures to employ a systematic branchegetic strategy that explores the impact of Ras interface mutants on binding to all effectors. Altogether, our results contribute to a quantitative and systems-level understanding of Ras-effector interactions and further our understanding of Ras in health and disease.

## Results

### Structural analysis of experimental complex structures

From the few experimentally determined complex structures of Ras with these RBDs, there are some structural similarities that can be determined (Figure 1). The main interface on the site of Ras consists of the two switch regions, switch 1 and switch 2. One of the main structural features of the interface between Ras and RBDs is the formation of an intra-molecular β-sheet between β2 on RAS and β2 on the RBD. This interaction is a highly conserved structural feature across all available complex structures, with deviations in orientation of less than 1 Å (Figure 1B, see Methods for MAE).

Analyzing the energetic profile of the interface *in silico* using the FoldX force field shows that there are also some recurring hotspots in the interface (highlighted in Figure 1C). I36, D38 and Y40 are well characterized as important residues for the interaction between Ras and effector domains. Additionally, for the structures 3DDC and 1LFD, the mutation E31K was introduced to stabilize the complex for crystallization. Our analysis confirms that this interface mutation has indeed been favorable for the complex formation. Finally, the energetic contributions of the function regions switch 1 and 2 to the interface were analyzed. Both relative and absolute contributions are diverse, although switch 1 contributions to binding dominate (Figure 1D). Altogether, the analysis of existing Ras-RBD effector structures indicates that although there are many common features of the Ras RBD interface such as the intra-molecular β-sheet or the hydrophobic patch around I36, the actual energetic contributions can come from different parts of the interface. It also sets the basis for a successful homology modelling approach.

### Homology modelling and characterization of modelled interfaces

In order to study these interface features in a more diverse set of structures, homology models of the complex between RAS and RBD were constructed for all proteins containing an RBD in the human proteome.

The homology modelling pipeline is based on the already existing complex models (Table S1). Also, with the recent release of AlphaFold2 ^14^ and the accompanying AlphaFold Protein Structure Database ^15^, the RBD domains of all potential effectors were extracted from that database and used. The structures of RBDs are predicted with good confidence by AlphaFold2 and our analysis indicates that AlphaFold2 is reliable at predicting the RBD fold (Figure S1). Additionally, AlphaFold2 complex modelling was attempted for all potential complex structures and models which AlphaFold2 were confident in (by Predicted Alignment Error (PAE)) and where the β-sheet alignment of the interface was within a tolerance to what has been observed in crystal structures, were used as templates as well (Figure S2 and S3, compare MAE Figure 1). There are two kinds of templates to use: 1) complex templates which comprised of the already experimentally determined complex structures of Ras and RBDs, as well as “good” AlphaFold2 predicted complex templates (Table S1), and 2) templates of the RBDs alone which were extracted from the AlphaFold2 Protein Structure Database (Table S2). An overview over the pipeline is depicted in Figure 2. Homology modelling was performed using homelette ^22^ with modeller and altMod ^23–25^. Evaluation of predicted structures was performed using QMEAN, MolProbity and SOAP potentials ^26-29^. The top 300 models for each target for each source of complex templates (experimental or AlphaFold2) were selected by combining the different scores. Subsequently, analysis using FoldX (interaction energy and alanine scan) was performed.

**Figure 2.**
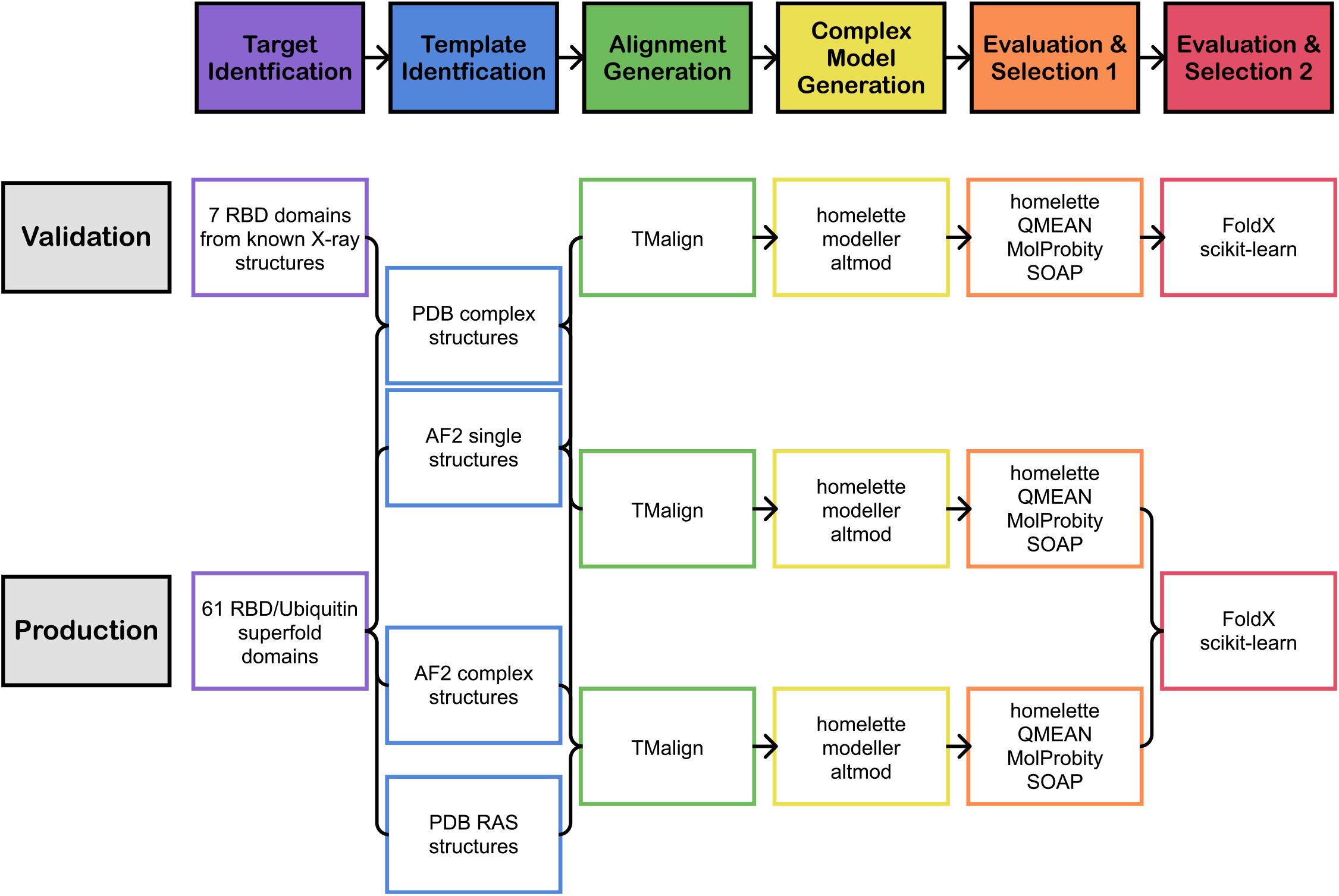
Overview over the homology modelling pipeline. The principal homology modelling steps are shown from left to right with the steps of “Target Identification”, “Template Identification”, “Alignment Generation”, “Complex Model Generation”, “Evaluation & Selection 1”, and “Evaluation & Selection 2”. In the “Validation” process, the homology modelling pipeline is run on a set of 7 RBD domains from known x-ray structures. The “Production” process describes the generation of 61 domains (RBD and Ubiquitin super-fold domains as controls).

In order to evaluate our approach, we generated a validation set, in which we created models for the structures already solved by X-ray crystallography without using information from the specific structure. Models generated in this validation set were compared to the underlying ground truth by assessing their correlation of the *in silico* alanine scan results to those of the crystal structures of interest. Using this ground truth, different methods to select representative models from the hundreds of structures were assessed. In particular, we evaluated the hyperparameters of an unsupervised learning pipeline comprised of different feature selection and dimensionality reduction strategies followed by clustering with the OPTICS algorithm (see Methods for more details about the hyperparameter space). After clustering, three representative structures were chosen. Based on the performance in the validation set, the optimal set of hyperparameters was chosen (Table S3, see Figures S4). The described strategy for identifying representative structures was then applied to all target complexes and we ended up with three representative complex structures for each effector.

Analogous to how we characterized the interfaces of the experimentally determined complex structures before, we performed the same analysis on the complex models. The overall FoldX interaction energies for the models are diverse, indicating that maybe some of the complexes are energetically unfavorable and would not form (Figure S5B). In general, the binding energies are lower than what would be observed in crystal structures, which is to be expected. There are one or two outliers with regards to FoldX binding energy, namely RASSF8 and PIK3C2B. In particular, RASSF8 is also showing an uncommon hotspot profile, with multiple unfavorable hotspots that are only appearing for this set of structures (Figure S5A). Based on this behavior, RASSF8 is excluded from further analysis.

The analysis of hotspots confirmed the already established hotspots. I36, Y40, D38 and E37 are the most commonly observed hotspots (Figure 3AB, Figure S5A). Interestingly, while I36, Y40, and E37 are exclusively favorable to the interaction with the effector protein, D38 seems to be also unfavorable in some of the structures (Figure 3B). The energetic diversity of the hotspot D38 was further investigated in the models. For this, two models were picked for which D38 was a favorable hotspot in the alanine scan analysis (Figure 3C: RASSF1, Figure 3D: RGL3), and two models were picked for which it was unfavorable (Figure 3E: ARAP2, Figure 3F: RAPGEF3). Next, neighboring amino acids were analyzed. For favorable interactions, we were able to observe positively charged amino acids. On RASSF1, we find K216 and H217, whereas on RGL3, there is R630. These amino acids probably form strong interactions with the negatively charged D38 on Ras. In contrast, for the models where D38 comes up as an unfavorable hotspot in the interface, we observe an uncharged, mostly hydrophobic neighborhood.

**Figure 3.**
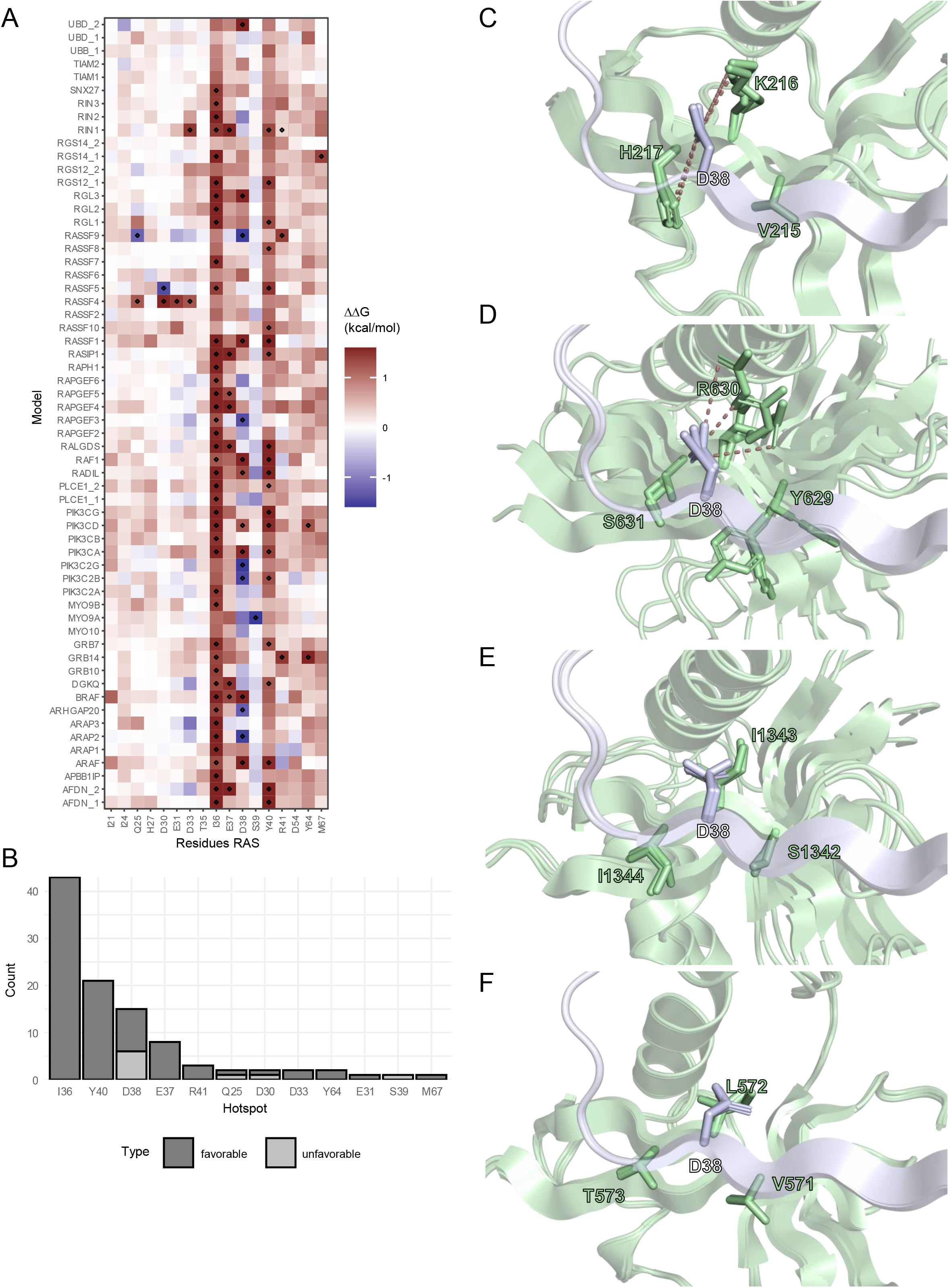
Energy hotspots in modelled structures. (A) Heatmap of FoldX alanine scan averaged from the three representative structures on each target. The color scale is confined to the limits [-1.6, 1.6]. Hotspot residues with a ΔΔG >= 1.2 or <= −1.2 kcal/mol were marked. Positive and negative ΔΔG values indicate a destabilization and stabilization of the RAS effector models by alanine mutation at the indicated position, respectively. C-F) Local neighborhood of D38 in structures where D38 is a favorable hotspot (C: RASSF1, D: RGL3) or an unfavorable hotspot (E: ARAP2, F: RAPGEF3). Ras and the effector structures are visualized in blue and green, respectively. Polar interactions between charged amino acids are indicated with dashes.

### Estimation of binding energies

One of the applications of the structural models that were generated was to use them for the estimation of binding energies. Since experiments measuring the binding energy between two proteins are experimentally very challenging and error-prone ^30^, we were implementing an *in silico* approach. Also, while FoldX is good at predicting energy changes to interaction energy or protein stability on mutation, the absolute interaction energies for protein complexes usually are not well correlated with experimental values ^31^. Because of this, a supervised learning regression pipeline was built based on features extracted from the modelled structures and a collection of experimentally determined binding energies of different Ras-effector complexes (Table S5). From a combination of different regressors, feature selection procedures, and hyperparameters, the best approach was determined using a cross validation strategy (see methods). A support vector machine-based regressor (see hyperparameters in Table 4) performed best in cross-validation with an R2 score of 0.53. Then, the performance of the best approach was evaluated in an out-of-sample test set, where it achieved an R2 score of 0.77. The model was then used to predict the interaction energies for the complexes without prior experimental measurements (Figure 4, Table S5). The predicted binding affinities for our models range 4 orders of magnitude, between the highest predicted affinity for BRAF of 0.02 μM to PIK3C2B with the lowest predicted affinity of 588 μM. The highest binding effectors are quite well characterized (RAF family, PI3K, RASSF5, RIN1, RalGDS, AFDN ^8^). As experimental measurements of the PI3K family members are difficult as the RBD is not easy to express and purify in isolation, it is noteworthy that we assign three of the good binding affinities to PI3K family members (PIK3CA, PIK3CD, PIK3CG). A big group of effectors has affinities in the range of 1 to 10 μM. For example, RASIP1 was previously in the “likely no binding” category ^8^, and is now predicted to have an affinity in complex with Ras of 2.7 μM. RAPGEF5 and RGL3 were previously in the “unknown” ^8^ category and have predicted affinities of 9.2 and 5.6 μM, respectively. Another big group of effectors has affinities in the range of 10 to 100 μM. Especially the first one could be interesting for modulation of binding affinity by the piggy back mechanism ^7^.

**Figure 4.**
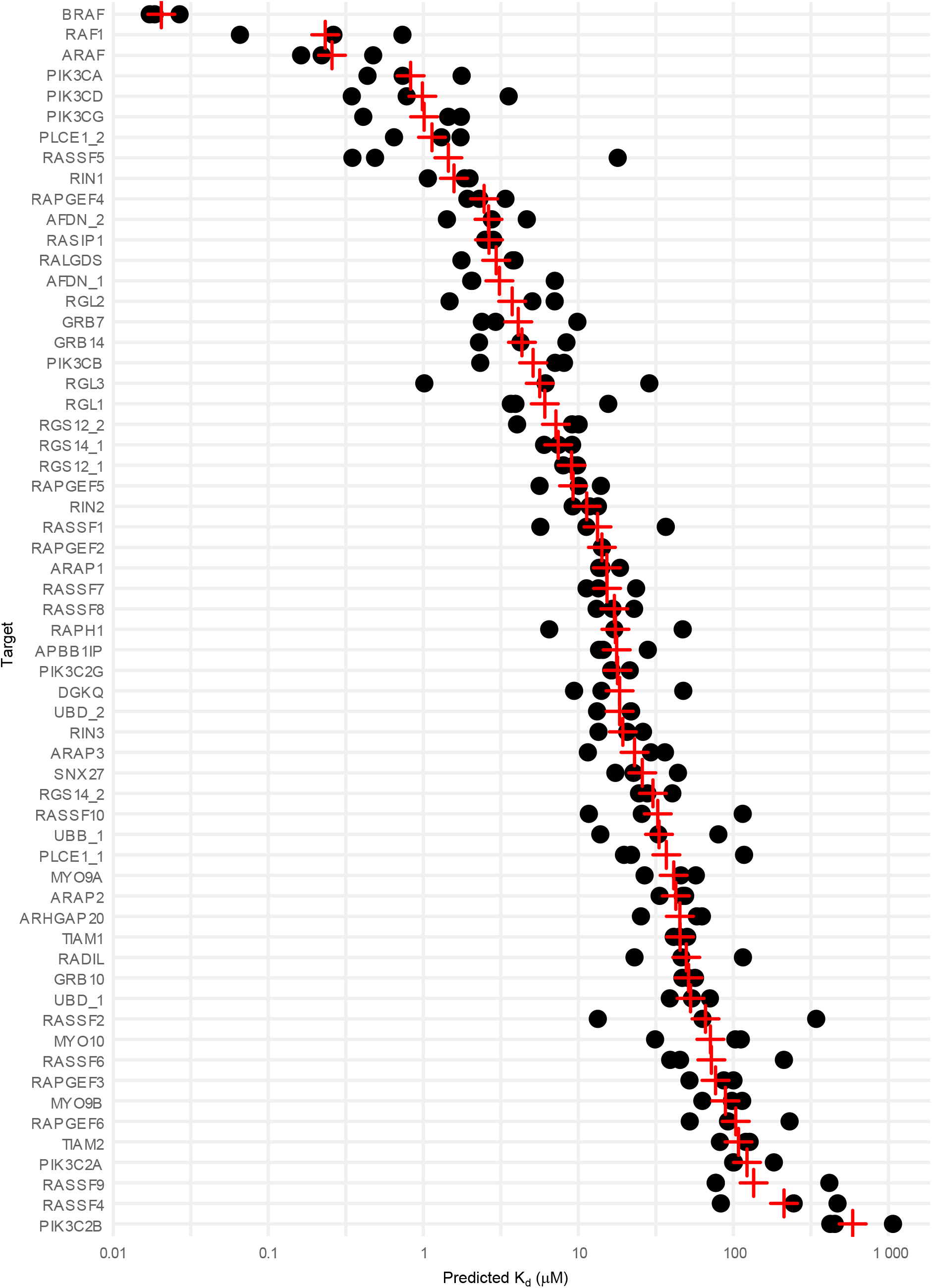
Results of the affinity prediction for all Ras-effector complex structures. Visualization of predicted binding affinities. The three representative structures for each target are visualized as black dots, with the averaged affinity (based on averaged energy) is visualized as a red cross.

### Switch contributions to binding

Having Ras-effector structural models available allowed us to analyze the individual contributions of switch 1 and switch 2 to binding using the results from alanine scanning (similarly as done before for the X-ray Ras-effector structures). We find that generally most structures are dominated by favorable switch 1 contributions (Figure 5 and Figure S6). Switch 2 contributions are surprisingly small. We also predict more contributions from regions outside switch 1 and switch 2 as in the X-ray structures. For the two weak affinity binding groups (10 μM (K_d_ < 100 μM and (K_d_ > 100 μM) switch 1 contributions are in the range of 4 to 5 kcal/mol with small (~1 kcal/mol) contributions from switch 2 and remaining parts involved in interface formation. For the two strong affinity groups (K_d_ < 1 μM and 1 μM (K_d_ < 10 μM) switch 1 contributions increase to 6 to 9 kcal/mol with also increasing switch 2 contributions (1 to 2 kcal/mol). We also observed negative switch contributions (mainly for switch 1), indicating that these proteins are either not well predicted or non-binders. Indeed, all structures with negative switch energies are predicted to be weak binders.

**Figure 5.**
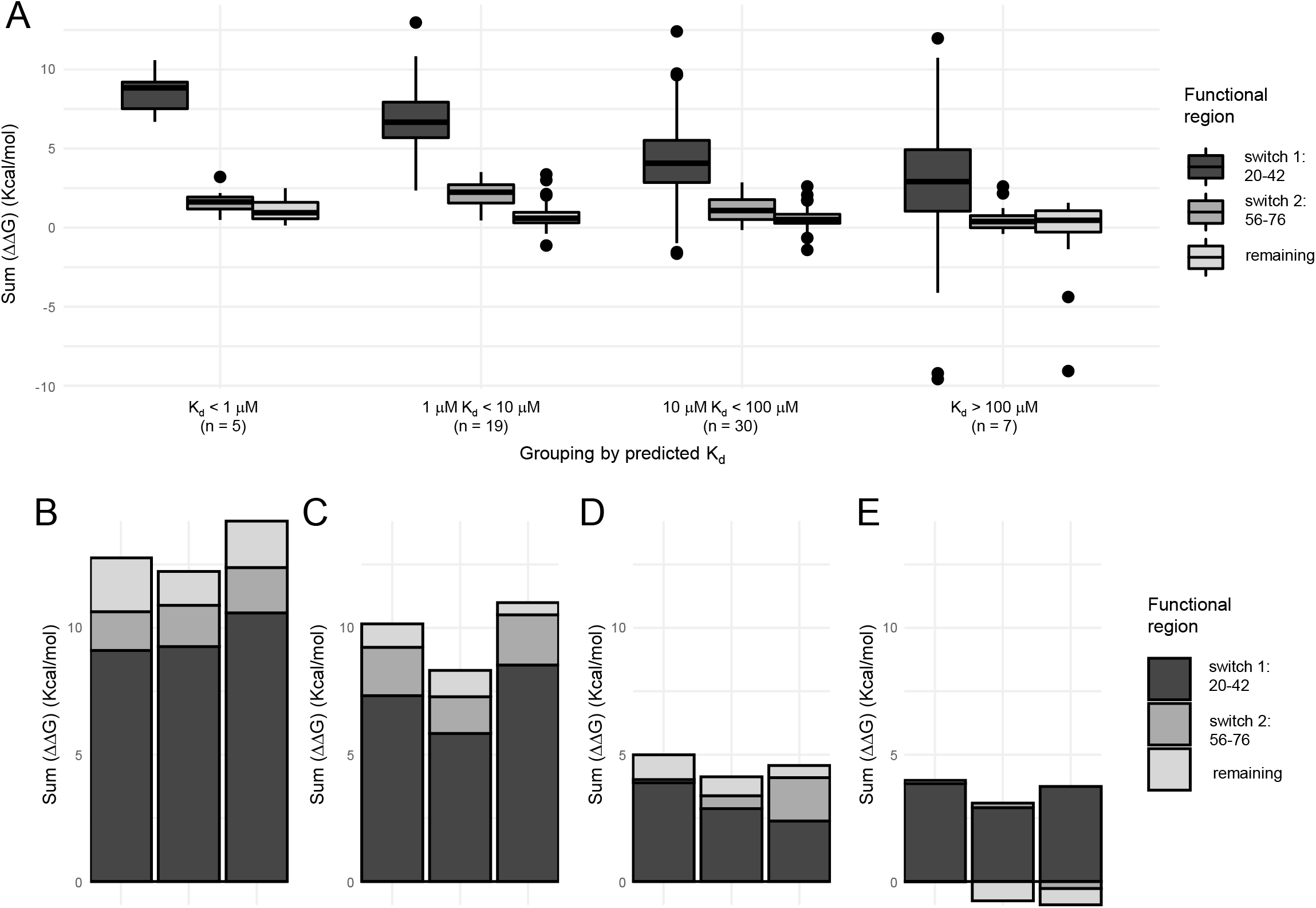
Switch contributions for the summed-up energy contributions grouped by their predicted binding affinity. (A) Energy contributions were separately calculated for switch 1 switch or the rest of the Ras interface by summing up energy energies from the *in silico* alanine scan analysis. Complexes were grouped based on their predicted binding affinities into four groups. (B – E) Examples of the switch contributions for each of the four groups. Visualized are BRAF (panel B), AFDN_1 (panel C), ARHGAP20 (panel D), and PIK3C2B (panel E).

### Branch pruning analysis using Ras-effector model structures

Next, we were interested in exploring surface mutations on Ras that would selectively influence the binding to some, but not all effectors. Both enhancing and inhibiting mutations are of interest. This could enable the engineering of the Ras effector system to respond to stimuli in different ways and to study selective sets of effectors. We previously reported a framework for the identification and evaluation of so called ‘branch pruning’ mutations ^21^. Since our protein is interacting with a many different effectors at the same time through the same interface it will be quite unlikely to identify mutations that selectively target only one protein. Instead, it is more likely that we will identify mutations that enhance in interaction with some proteins while inhibiting some others.

Figure 6 shows a heatmap of all identified mutations of interest and their effect on all effectors. Some interesting mutations to highlight are mutations around I36, that are almost exclusively unfavorable while affecting almost every structure. D37 mutation are more selective and also exclusively disruptive. D38 is mixed, as our analysis of the hotspot already indicated. This is probably the best point to disrupt the system. Y40, interestingly, while being a ubiquitous hotspot, is not a good spot for engineering the interface because mutations seem to affect protein stability (compare with Figure S7A). Also of interest is that we detect both mutations that increase and disrupt binding (Figure S7B).

**Figure 6.**
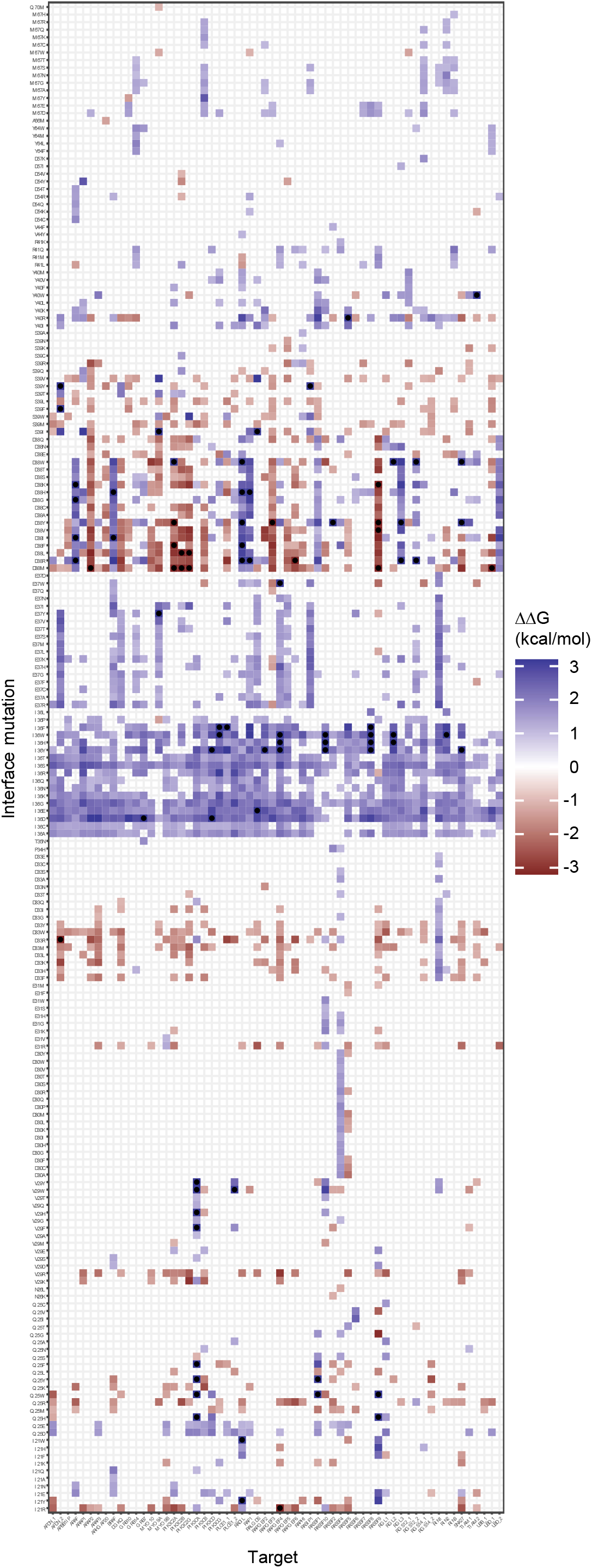
Energetic characterization of RAS interface mutations that affect effector binding. The color scale is confined to the limits [-3.2, 3.2]. Hotspot residues with a ΔΔG >= 1 or <= −1 were marked. Negative ΔΔG values indicate an increase in binding energy compared to the WT interface, and positive ΔΔG values indicate a decrease in binding energy compared to the WT interface.

### Assessment of rewiring of Ras-effector interaction on a systems level

Finally, we want to evaluate the behavior of the Ras effector system based on our predicted binding energies and based on the introduction of different branch pruning mutations. For this means, we went back to our mathematical model of the Ras-effector system that incorporated affinities and high-quality proteomics data in 29 human tissues ^7^. Here, all 29 tissue systems were simulated at 20% and 90% GTP load on Ras. We are using exclusively the predicted binding energies for this (except RASSF8, see methods). Overall, the results for the systems without a branch pruning mutation are comparable to previous findings (Figure 7A; ^7^). The Raf family members ARAF, BRAF and RAF1 dominate the binding profile in complex with Ras. Other effectors that are in high amount in complex with Ras in at least one of the 29 tissues are RGL2, RASSF7, RASSF5, RASIP1, RALGDS, PIK3CD, PIK3CA, and AFDN.

**Figure 7.**
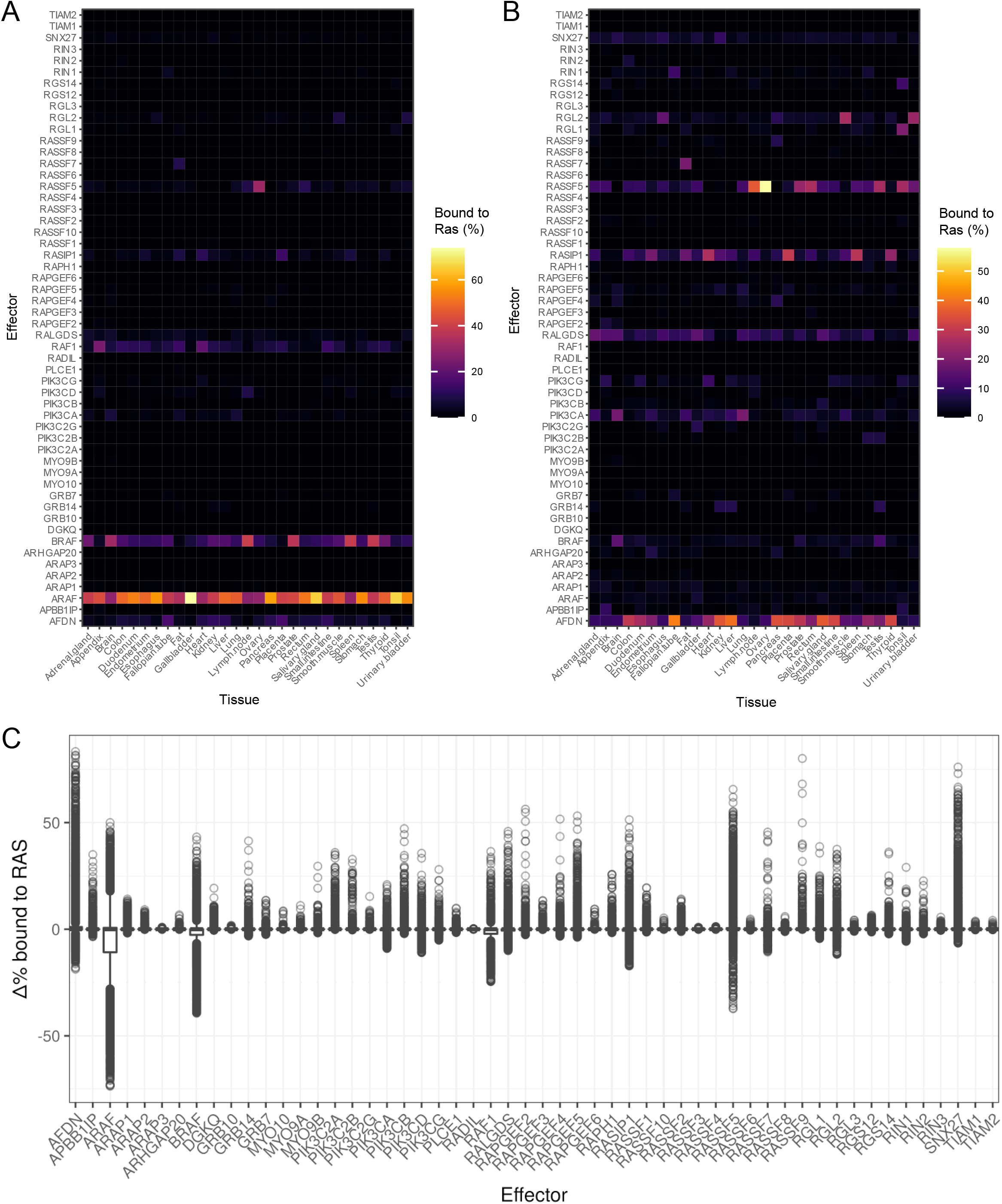
Ras-effector interaction rewiring at a systems level. (A-B) Effects of interface mutation on the Ras-effector system. Heatmap of effectors bound to Ras at 20% Ras-GTP load in 29 tissues in (panel A) WT interface and in (panel B) with the effects of a D38A mutation. (C) Visualization of change in relative binding compared to WT system for each effector across all tested branch pruning mutations in all 29 tissues at 20 % or 90 % Ras-GTP load. Positive values indicate a relative increase in bound effector to Ras.

Expanding on this, we introduced our branch pruning mutations to the mathematical model. In total, there are 200 interface mutations that do not significantly affect overall protein stability but affect binding to at least one of the effectors. Some of the branch pruning mutations are able to dramatically change the system in all tissues, as can be seen from the example of D38A (Figure 7B). With a single interface mutation, almost all RAF binding is quenched, and other effectors start to compete for the binding. Interestingly, which effectors come up depends on the tissue.

Next, we explored to which extend, for a specific effector, it is possible to modulate its binding. For this, we visualized the possible changes to the relative amount of effector bound to Ras across all systems tested (Figure 7C). On the side of proteins that can be negatively influenced, mostly the high-affinity binders such as RAF family proteins show up, which is to be expected. Some effectors cannot be influenced by the branch pruning mutations, either because they are energetically not affected or because their concentrations in any of the 29 tissues do not leave them in a position to compete for binding. Examples for this would be RADIL or the TIAM family proteins. The effectors with the highest propensity to have their binding enhanced are AFDN, RADIL, SNX27, RASSF5.

Finally, we investigated whether there are recurring states that the modelled system assumes and whether these states are dependent on Ras-GTP load, the tissue, or interface mutation. To this end, we applied uniform manifold approximation and projection (UMAP) to our systems to transform a high-dimensional space of absolute and relative effector binding into a 2D space. Then, we used OPTICS to identify areas of high density in this 2D plane and assigned them into 24 distinct clusters as well as outliers outside the clusters (Figure 8A). For each cluster, we picked three of the systems at random and visualized their relative effector binding (Figure S8). Each of these clusters belongs to a different “state” that the Ras-effector system can be rewired into, with systems belonging to a specific state showing similar trends in effector binding. Many of the systems are dominated by ARAF binding, as it would be expected, but even for these there are differences in the secondary effectors. To understand what the attributes of different “states” of the systems are, two of them were picked and investigated for Ras-GTP load, tissue composition, and interface mutation status (Figure 8BC). We find that there are different ways a distinct state of the system can come together: for the state analyzed in Figure 8B, we can see that it is composed of many different tissues, but only a handful of interface mutations, most of them D38 mutations. This indicates that this state can be reached from many different tissues by a specific, recurring set of mutations. In contrast, the state analyzed in Figure 8C is entirely composed of a single tissue (lymph node) with many different mutations, indicated that this state can only be reached by a specific tissue. Interestingly, both states analyzed are diverse in terms of Ras-GTP load. To conclude, we identify 24 distinct states of the Ras-effector systems and show that there are different mechanisms on how these states are formed.

**Figure 8.**
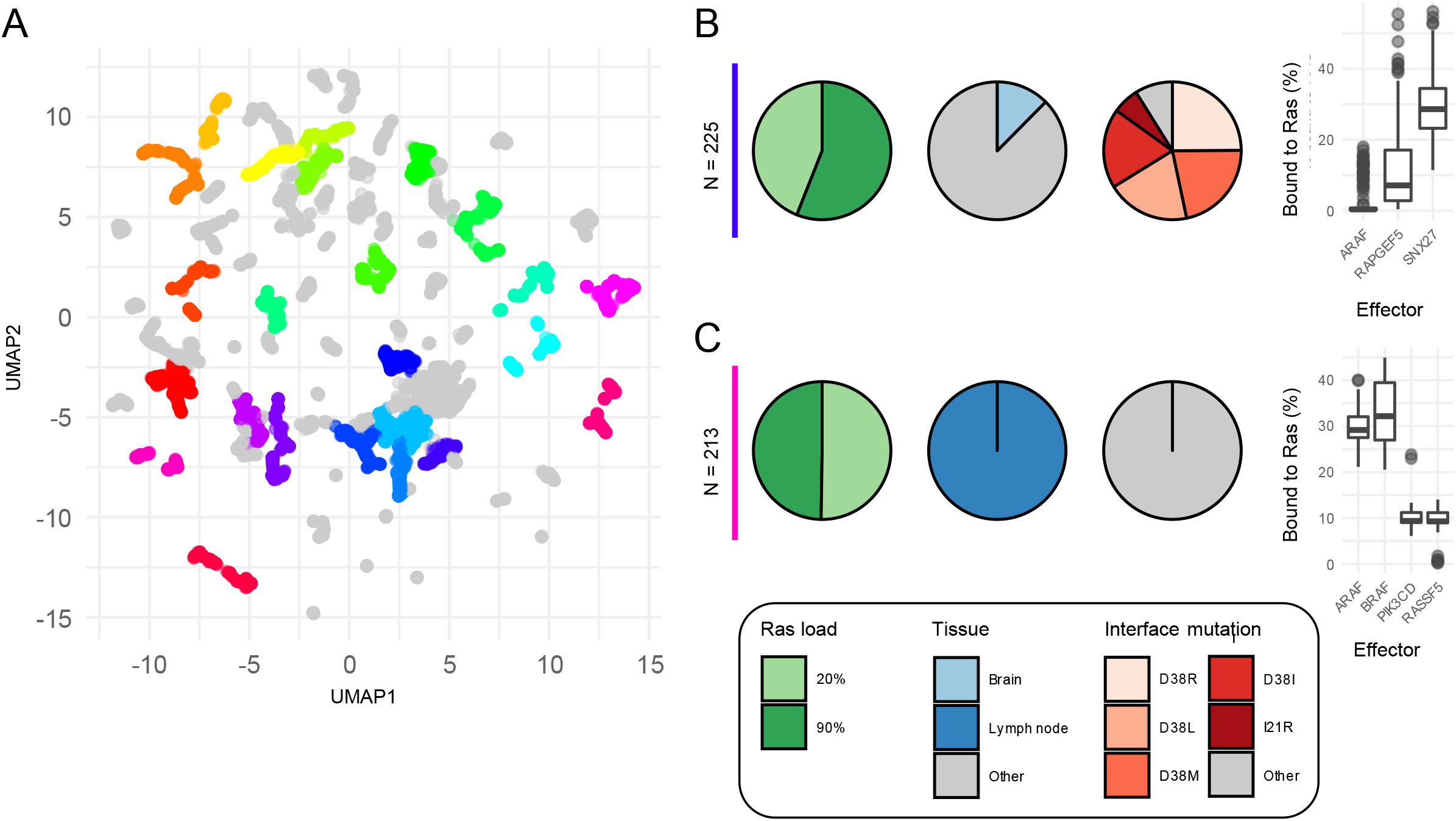
Ras-effector systems are rewired into distinct states. (A) UMAP transformation of all simulated systems. Similar systems were identified using OPTICS clustering. 24 identified clusters are colored in rainbow colors, with systems classified as outliers colored in grey. (B) and (C) Characterization of two identified state clusters in terms of Ras-GTP load, tissues, interface mutations, and most representative effectors. For the tissue and mutation pie charts, all groups that were smaller than 5% of the total observations were collapsed into the “Other” group.

In summary, based on a large set of Ras-effector models, we predicted Ras branchegetic mutations and evaluate their binding in a tissue-specific Ras competition model. The Ras mutations, once introduced into cells or tissues, can be used as a tool to probe the contribution of specific effector pathways to an output or cellular phenotype.

## Discussion

In this work we have shown a complete structural reconstruction of Ras and the RBDs of its effectors based on state-of-the-art technology. We used these structural models to investigate the workings of the interface between Ras and its effectors, as well as to search for and identify potential branch pruning mutations on Ras that would alter the behavior of the underlying system. We analyzed the effects on the steady state of the Ras effector system.

Recent advances in structural modelling, mostly by the development and release of AlphaFold2 and similar algorithms have pushed the field of structural bioinformatics forward. This development was crucial for the quality of this study. While a normal homology modelling pipeline would have been successful, the inclusion of both aspects of AlphaFold2 models was essential for the quality of the results. On the one hand, finding additional potential complex templates diversified the possible configurations of the interface that we could use to generate our models. On the other hand, having high quality templates of the RBDs enabled us to improve the structural predictions. Interestingly, with all the advancements that AlphaFold2 brought, this combined AlphaFold2/homology modelling approach yielded more consistent and better results for this particular question.

There is a growing body of literature about how Ras dimerizes or forms multimers, or interacts with the membrane to modulate the signaling. All these aspects have been deliberately left out for this approach. The essence of the interaction of Ras and its effectors is the binary protein-protein interaction between the Ras switch regions and the RBD. This common feature was the focus of this work, and we believe that other factors such as dimer/multimerization of Ras, the composition of the membrane, etc., are only modulators for this interaction.

By creating a complete structural system, we were able to investigate and understand the interactions of Ras with its effector proteins on a different level. Crucially, it enabled us to analyze how *in silico* mutations of the system could affect its behavior. This is an interesting approach for a lot of different systems, not just the Ras effector system. However, there are certainly challenges. The prediction and verification of a protein-protein interface can be very complicated, and the techniques for modelling protein-protein interactions are not sufficiently developed to easily translate that approach to a larger scale. Probably the main reason why it was possible to construct this structural system for the Ras-effector system was because it is a conserved domain-domain interaction between homologue proteins. Although the sequence identity of the RBD sequences is not well preserved anymore, the structural fold of these domains has been preserved. Additionally, there is also the preserved mode of binding by the formation of the intermolecular β-sheet. These factors were favorable for the construction of the structural system and would need to be addressed if this approach were to be taken to a higher scale. Recent publications are already able to work on system-wide structural prediction for interactions ^32^. These approaches are very promising for the structural characterization of known protein complexes and can identify high-confidence novel interactions as well. However, for the exhaustive characterization of a full system, especially with transient interactions, the distinction between true and false positives seems to remain challenging.

Finally, we hope that our structural and systems analysis of the branch pruning interface mutations will enable interesting experimental setups that study different downstream pathways from Ras. Some of the more promising candidates are AFDN, RADIL, SNX27, RASSF5. The downstream pathways of Ras that are best characterized are the RAF-MEK-ERK signaling pathway and the PI3K-AKT signaling pathway. However, some of the other proteins might play a role in a physiological or pathophysiological context as well. AFDN is essential for the organization of adhesion between cells ^33^, function that is often impaired in cancer ^34^. RADIL is also linked to cell adhesion, and recent data showed that knockdown of was linked to decreased cell proliferation and invasion ^35^. SNX27 is also part of signaling pathways that link to cell adhesion and barrier function ^36^. RASSF5 is a tumor suppressor and has been shown to inhibit growth and invasion and to induce apoptosis ^37^. Importantly, all branchegetic mutations were studied on a systems level using a tissues specific Ras competition model. These models can easily be adapted for specific cell systems of interest, provided that estimates for Ras and effector abundances are available. Altogether, this work contributed to a quantitative understanding of a key cellular hub protein – Ras.

## Acknowledgements

The authors would like to thank Cian D’Arcy, Thomas Sevrin, Simona Catozzi, Swathi Ramachandra Upadhya, and Hiroaki Imoto for discussions and/or critical reading of the manuscript.

This work is part of the research program “Quantitative and systems analysis of (patho)physiological signaling networks” [16/FRL/3886], which is financed by Science Foundation Ireland (SFI) to C.K.

This publication has emanated from research conducted with the financial support of Science Foundation Ireland under Grant number [16/FRL/3886]. For the purpose of Open Access, the author has applied a CC BY public copyright licence to any Author Accepted Manuscript version arising from this submission.

## Author contributions

Conceptualization, P.J. and C.K.; Methodology, P.J. and C.K.; Software, P.J; Investigation, P.J.; Data Curation, P.J.; Writing – Original Draft, P.J. and C.K.; Writing – Review & Editing, P.J. and C.K.; Visualization, P.J. and C.K.; Supervision, C.K.; Funding Acquisition, C.K

## Declaration of interests

The authors declare no competing interests.

## STAR★Methods

### Lead contact

Further information and requests for resources should be directed to Philipp Junk and will be fulfilled by the lead contact, Philipp Junk

### Material availability

This study did not generate new unique reagents.

### Data and Code availability

The code and generated data have been deposited on Zenodo: https://doi.org/10.5281/zenodo.7188727

## METHOD DETAILS

### Experimentally determined Ras-effector complex structures

Structures were downloaded from the PDB. In the case where multiple models were available, the best one by MolProbity score was selected ^27 28^. The PDB files were processed using pdb-tools so that all GTP and Mg2+ annotations were in the expected format ^38^. The list of used template structures can be found in Table S1.

### Interface characterization

Hotspots residues were determined by FoldX *in silico* Alanine scan ^39 31 20 40 41^. In detail, each residue on both RAS and the effector was mutated to alanine and the *in silico* change of binding energy ΔΔG was determined as such:

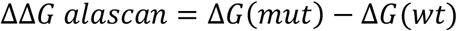

A positive ΔΔG value indicates that the mutated residue is involved favorably in the interaction, whereas a negative ΔΔG value indicates that the mutation to alanine improved the interaction between the two proteins. The standard error for ΔΔG values in FoldX is around +- 0.8 kcal/mol. In order to identify the most important residues for the respective interaction, a ΔΔG cut-off of 1.6 kcal/mol was chosen for the investigation of crystal structures. Since the energies are systematically lower in the modelled complex structures, a cut-off of 1.2 kcal/mol was chosen for those.

Based on the alanine scan results, the contribution of the two major functional regions in the RAS interface ^42^, switch 1 (residues 20-42) and switch 2 (residues 56-76) to the interaction were determined. For each functional region and the remainder of Ras, the ΔΔG values were filtered by abs(ΔΔG) > 0.8 kcal/mol and then summed up. The definition of the switch regions used here is more generous than what is normally used in the literature, and it would be probably more correct to label them as “switch-influenced” regions. These residue ranges aim to capture the regions in the interface that are affected by the movement of switch 1 and 2 during the transition from the inactive GDP bound state to the active GTP bound state. The conserved intra-molecular β-sheet between the β2 sheet on Ras with the β2 sheet on the effector is evaluated by measuring the differences in inter-molecular distances between the β sheets in the experimentally solved complex structures and comparing them to the ones in a structure of interest. This measurement has been named Measured Alignment Error (MAE) in this manuscript:

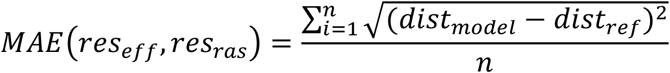

with *dist_model_* being the Euclidean distance between two residues in the model of interest; *dist_ref_* being the Euclidean distance between the residue on Ras and the corresponding residue in the reference structure, as determined by structural alignment using TMalign ^43^; and *n* being the number of reference structures. The references used were the X-ray complex templates found in the PDB (Table S1). When the MAE is calculated for X-ray complex templates, the comparison with itself is removed from the calculation.

### AlphaFold2 determined single structures for RBDs

For each protein containing one or more potential RBD sequences as identified in ^6^, the full structure was obtained from the AlphaFold Protein Structure database ^14 15^ (Release 1, accessed September 2021). The sequences of the RBDs were obtained from Pfam ^44^ and UNIPROT ^45^. The part of the structures that correspond to RBD sequences was subsequently extracted from the AlphaFold2 structures. The list of used templates as well as information about the extracted RBD domains can be found in Table S2.

### AlphaFold2 determined complex structures: generation and selection

AlphaFold2-multimer/ColabFold (version 1.3.0) was run on a local computer ^46 14 47^. Multiple sequence alignments were generated from MS2seq ^48 49 50 51^. Complex models were generated with HRAS as interaction partner. For each target, five models were generated with additional template search and five without additional template search ^46 52^. The resulting model were relaxed as per ColabFold defaults ^46 53^. For each model, several metrices were evaluated. Firstly, AlphaFold2’s Predicted Alignment Error (PAE) was taken into consideration. PAE is an estimation of the error of pairwise distances between residues, that can be used to assess how confident AlphaFold2 is in the inter-domain arrangement of its models. Secondly, FoldX ^39 31 40 41^ interaction energies were determined for the complexes as described above. Finally, the expected orientation of the inter-molecular β sheets was evaluated by MAE. The best model by MAE was selected for each target, and then all complex structures with a MAE > 1A were removed.

### Homology modelling pipeline

Alignment generation, model generation and model evaluation were performed using the homelette homology modelling interface ^22^. Inputs to the homology modelling pipeline were complex structures of Ras in complex with some effectors, either experimentally determined or selected from AlphaFold2 complex predictions, as well as AlphaFold2 models of all RBDs of interest. For each target, all combinations of the RBD template with all complex templates were used to generate 300 models of the target in complex with HRAS each. The different sources for the complex templates were run separately with slight differences in the modelling procedure. All sequence alignments for RAS were generated with M-Coffee ^54,55^. For complex templates of experimental origin, TMalign ^43^ was used to generate a structure-based sequence alignment based on the RBD in the complex template and the single RBD of the target structure. Then, with those two templates as inputs, models were generated. For complex templates of *in silico* origin, structure-based sequence alignments were generated with TMalign ^43^ as described. As an additional template a HRAS single structure (5P21 ^3^) was used in the modelling process since the AlphaFold2-generated complex templates, in contrast to the complex templates of experimental origin, do not have information about the important heteroatoms GTP and MG2+ in the Ras part of the structure. Models were generated using modeller ^23 24^ with the altMOD extension ^25^. All models generated were evaluated using QMEAN ^26 56^, MolProbity, ^27 28^ and SOAP ^29^ potentials. A combined score was determined based on borda count as such:

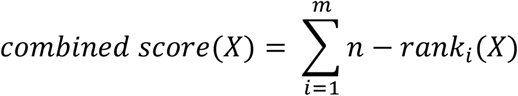

For an observation X with i… m being a collection of evaluation criteria and n the total number of observations. A structure with high borda score is a structure that performs well across all metrices.

For each of the different sources of complex templates (experimental or *in silico*), 300 models were selected in a first selection step based on the combined score. These 600 models per target were then further analyzed using FoldX. *In silico* interaction energies were determined and *in silico* alanine scan was performed (see Interface Characterization).

In addition to generating models for unknown targets, a set of validation models based on the experimentally solved models were generated as well. For each of the seven PDB complex structures, 300 models were generated. Inputs were restricted so that the structure to be modelled would not be used as a complex template, but only the remaining six experimentally derived complex templates. For each validation target, 300 models were selected as described above.

Using the results from the analysis with FoldX, representative structures were selected based on an unsupervised learning workflow. As the clustering algorithm of choice, Ordering Points To Identify the Clustering Structure (OPTICS) ^57^ was used, as implemented in scikit-learn ^58^. OPTICS is a density-based clustering method, which unlike the more popular k-means clustering does not require a manually set input of the number of clusters. Additionally, OPTICS is able to label data points as outliers. Several approaches to feature selection and/or feature engineering, hyperparameters of OPTICS, as well as methods for selecting representative structures from clusters were evaluated against the set of validation models (Table 3). To select the best combination of hyperparameters, for each combination, the *in silico* alanine scan results of the representative structures were correlated to the results of corresponding PDB structures, and the combination of hyperparameters with the most stable performance across all validation sets (by minimum z-score of the correlation against the PDB structure for all 7 validation sets) was chosen. The best combination of hyperparameters is highlighted in (Table 3).

### Estimation of binding energies

Supervised learning based on a number of FoldX-derived features was used in order to estimate binding energy of complex models. The features consisted of the FoldX interaction energy, the energy contribution of switch 1, 2 and the remainder of the Ras protein interface (see Interface characterization), and the ΔΔG values for hotspot residues on Ras (cut off 1.2 kcal/mol). All features were standardized and highly intercorrelated features were removed. Data preparation was performed in R.

The experimental measurements of the dissociation constant between Ras-effector complexes were collected from several publications (see Table S5, also available as supplementary data). Effectors that were experimentally determined as non-binders were removed from the data set due to uncertainty how to encode them with the varying technical limitations on detectable binding energies at the time of their publication. Also, the models generated for RASSF3 seem to be outliers with regards to FoldX interaction energy (see Figure S5B) and were therefore removed from the prediction. A test set was manually chosen from the available experimental measurements to cover the full spectrum of experimentally determined interaction energies. At the end, this gave us a training set with 51 observations (17 * 3 models) and a testing set with 12 observations (4 * 3 models).

Supervised learning was performed in scikit-learn ^58^ using different combinations of feature selection algorithms and regressors. Feature selection was performed using either f-regression or mutual information as implemented by scikit-learn. Regression was evaluated for different algorithms with various hyperparameter spaces (see Table S4). All combination of feature selection and regression were evaluated using Group-K-Fold cross validation within the training data, with k=5 and the groups corresponding to the three structural models chosen for each target. Models were evaluated using R2 score. The best performing combination was trained on the whole training data and evaluated against the test data. Finally, this model was used to predict the binding energies for the complex structures without experimental data.

### Branch pruning

FoldX was used to evaluate the effect of interface mutation on the binding energies in the modelled complex structures. As previously described, ^21^ the FoldX command PSSM was used to evaluate the changes in binding energy, and the FoldX command PosScan was used to evaluate if mutations impacted the stability of RAS.

All mutations that were destabilizing HRAS in either of two structures (3TGP and 5P21) were removed from the analysis (cutoff 1.6 kcal/mol). Then, between the three models for each target structures, it was checked if a mutation had a noticeable impact (> +- 1kcal/mol) in at least two of the three structures. If so, the changes in binding energy for all models above/below the cutoff were averaged.

### Systems analysis

A mathematical model of the Ras-effector system was set up as previously described ^6 7^ The model is based on classic ligand-receptor kinetics according to the assumption of conservation of mass. A system of ordinary differential equations was set up and steady states were calculated as described. The reactions are expressed as such:

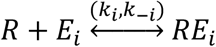

With *R* representing the molar concentration of Ras, *E_i_* the molar concentration of an effector of the set of *i* ∈ (1, 2, …, 54) effectors (all modelled proteins, except for proteins from the Ubiquitin family) and *RE_i_* the molar concentration of a Ras-effector complexes. The complex is formed at rate *k_i_*, and dismantled at rate *k_−i_*. These rates define the dissociation constant:

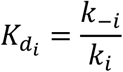

Due to the assumption of mass conservation,

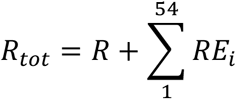

and

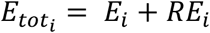

the system to solve therefore is:

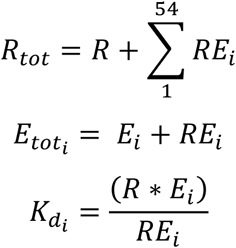

For the set of *i* = {1,2, …,54} and can be numerically solved for *RE_i_*. The system was solved using SciPy ^59^.

The concentrations for the species in the model were taken from the supplementary data of ^7^, in which molar concentrations were derived from high-quality proteomics data set of 29 different human tissues ^60^. The concentration of Ras proteins (HRAS, NRAS, KRAS) were pooled together and then multiplied with a loading factor to take into account the balance between active (GTP bound) and inactive (GDP bound) Ras. This loading factor was set to 0.2 for a normal RAS system, and 0.9 for a system hyperactivated by an oncogenic hotspot mutation.

The binding affinities were taken from the predicted binding affinities (see Estimation of binding affinities), with the exception of RASSF3 for which we are not confident in our structural predictions. An experimentally determined binding affinity for RASSF3 was substituted from ^61^. Additional sets of binding affinities based on the branch pruning analysis were evaluated as well. For this, the predicted binding affinities were adapted by the ΔΔG values from the branch pruning analysis. For the case that a system with hyperactivated Ras was considered, it was made sure that common oncogenic hotspots such as G12, G13 and Q61 do not influence the binding energies for any effector.

To gain an overview over all solved system, absolute and relative effector bindings were transformed using UMAP ^62^ and visualized. OPTICS clustering ^57^ was performed on the UMAP transformed data (parameters: min_sample = 25, min_cluster_size=200).

### Data analysis and visualization

All data analysis, unless otherwise noted was performed in R using the tidyverse environment ^63,64^. Visualizations were generated using ggplot2 ^65^. Visualizations of protein structures were generated using PyMol ^66^.

## Notes

https://doi.org/10.5281/zenodo.7188727

### Competing Interest Statement

The authors have declared no competing interest.

### Summary of Updates

Reorganized main and supplementary figures; updated Section "Assessment of rewiring of Ras-effector interaction on a systems level"; updated references

