## Supplementary Figures for "Structure-based prediction of Ras-effector binding affinities and design of ‘branchegetic’ interface mutations"

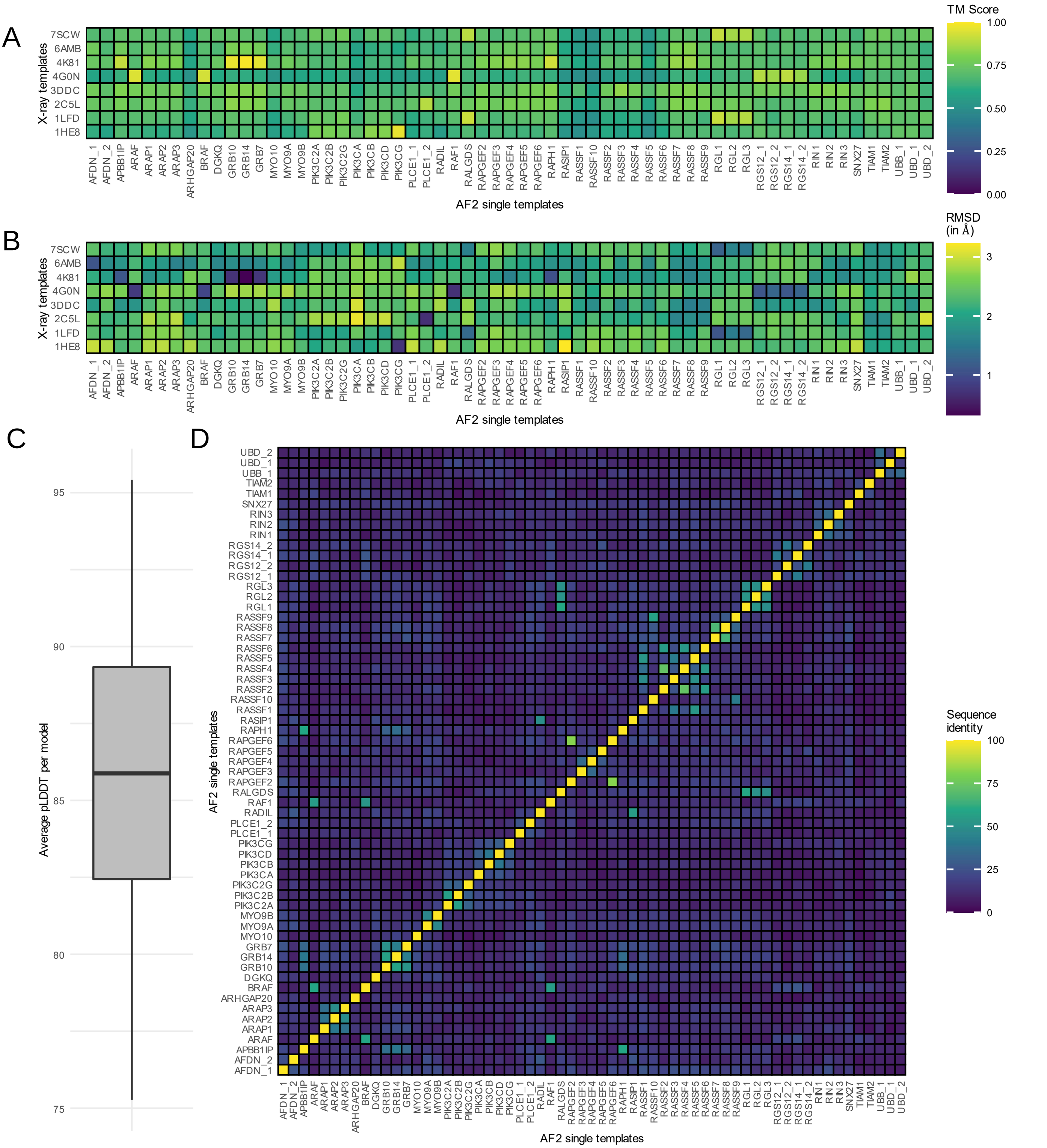


**Figure S1. Characterization and evaluation of AF2 RBD single templates retrieved from the AlphaFold Protein Structure Database**

(A) TM Score from the TMalign structural alignment program of AF2 generated RBD structures in comparison with the RBD domains of known complex structures. A TM Score between 0 and 0.3 corresponds to a random fold, whereas a TM Score of > 0.5 indicates a similar fold. The TM score shown is normalised to the length of the AF2 template.

(B) RMSD (in A) of AF2 generated RBD structures compared to crystal structure references. RMSD was calculated using TMalign.

(C) Distribution of average pLDDT values from the models retrieved from the AlphaFold Protein Structure Database. The pLDDT score is a local measure of how confidence estimated by AlphaFold2 when generating models. pLDDT scores > 70 and >90 correspond to confident and very confident predictions, respectively.

(D) Matrix of pairwise sequence similarity (in %) for all RBD/Ubiquitin-superfold domains investigated in this study.

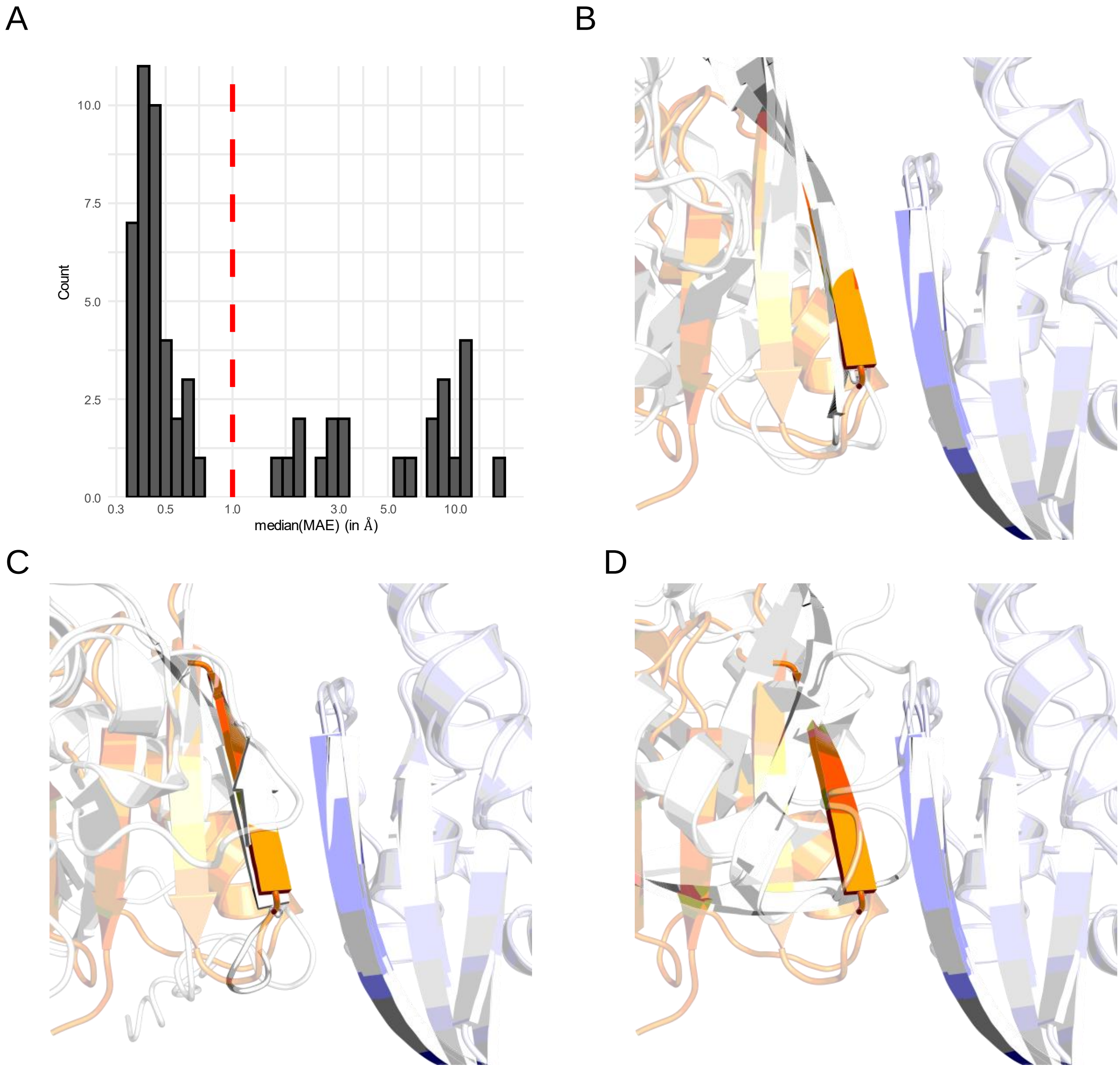


**Figure S2. Selection of AF2-generated complex templates by MAE**

(A) Histogram of the MAE distribution for the best structure for each target by MAE. The selection cut off was set to 1 Å. The x-axis is visualized in log10 transformation.

(B) Visualization of β-sheet alignment for complex models generated with AlphaFold2 which have low median MAE value (GRB10: 0.35, RALGDS: 0.41).

(C) Visualization of β-sheet alignment for complex models generated with AlphaFold2 which have moderate median MAE value (MY09A: 0.62, RASSF6: 0.68).

(D) Visualization of β-sheet alignment for complex models generated with AlphaFold2 which have high median MAE value (RASSF7: 1.53, MYO10: 1.77). In B, C and D, Ras is displayed in light blue, RAF1 from the X-ray structure 4G0N is displayed in orange and the structural models generated with AlphaFold2 are displayed in white. The proteins were made semi-transparent except for β2 on RAS and β2 on the effectors.

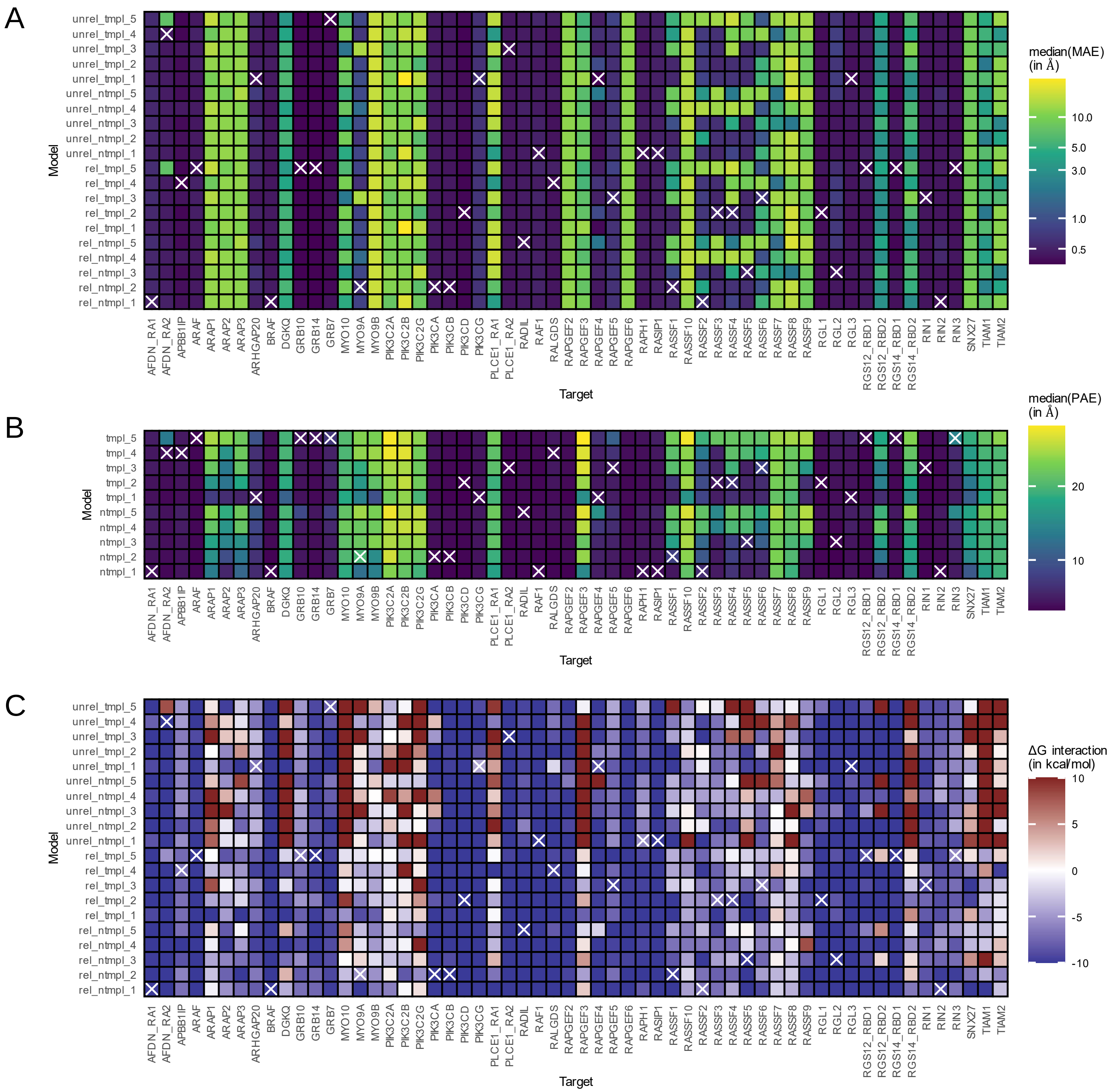


**Figure S3. Characterization of AF2-generated complex templates**

(A) Median Measured Alignment Error (β-sheet alignment) for AF2 determined complex structures in Angstroms. Selected structures are marked with a white cross.

(B) Median Intra-molecular Predicted Alignment Error for all AF2 determined complex structures. For the calculation of the median, only PAE values between the two chains in the model were considered, not within the two chains. Selected structures are marked with a white cross.

(C) FoldX interaction energy for all AF2 determined complex structures. Selected structures are marked with a white cross.

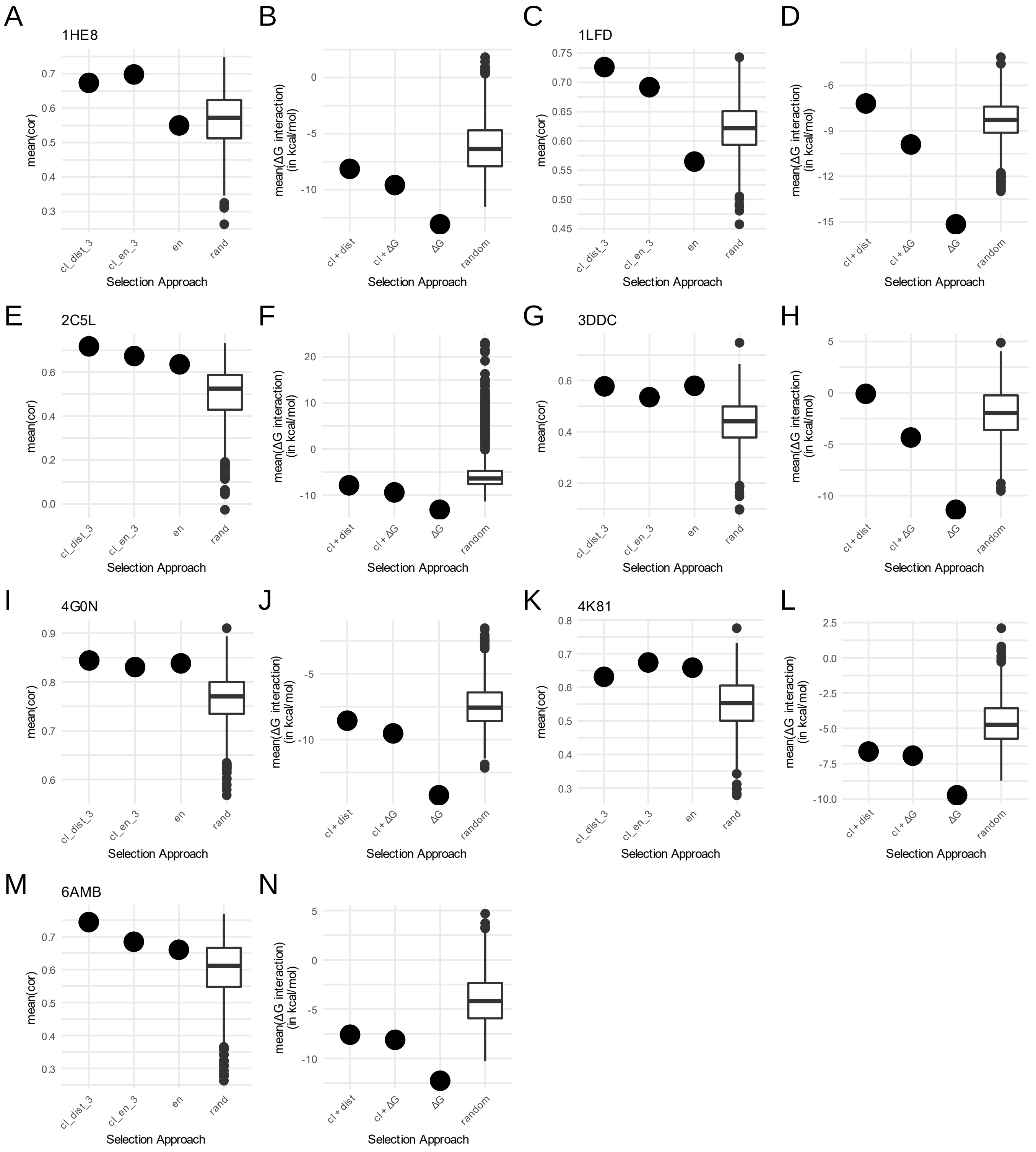


**Figure S4. Comparison of different strategies to select three representative structures for the validation set (A and B: 1HE8; C and D: 1LFD; E and F: 2C5L; G and H: 3DDC; I and J: 4G0N; K and L: 4K81; M and N: 6AMB).** Visualization of the average correlation of the selection (A, C, E, G, I, K and M) and the average FoldX interaction energy (B, D, F, H, J, L and N). Selection strategy cl+dist is the selection of three representative structures after clustering from the best cluster by FoldX interaction energy based on lowest average distance in the cluster. Selection strategy cl+ΔG is the selection of three representative structures after clustering from the best cluster by FoldX interaction energy based on the best FoldX interaction energy. Selection strategy ΔG is the selection of three representative structures without clustering based on the best FoldX interaction energy. Multiple draws of three random structures are visualized as a comparison.

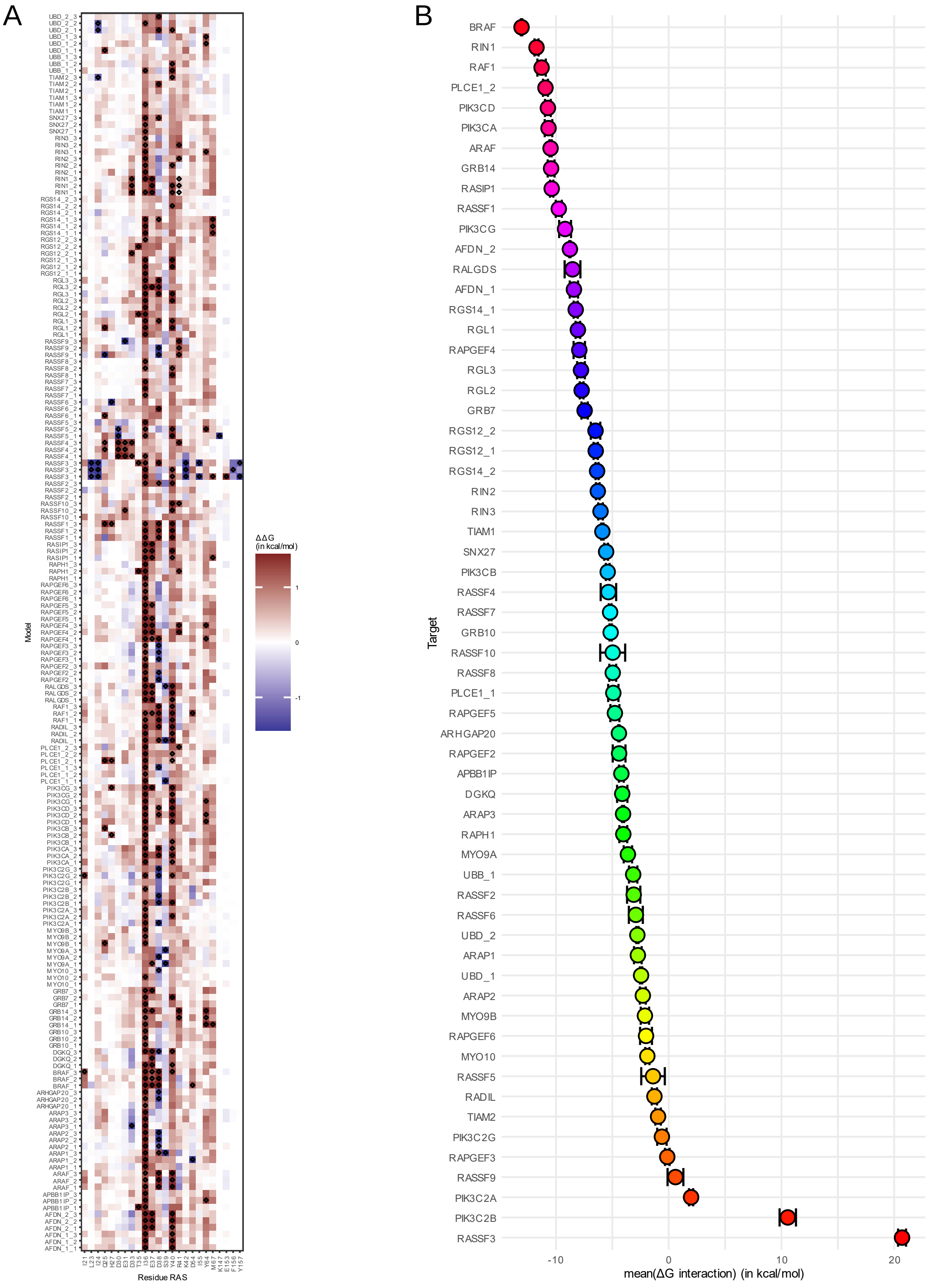


**Figure S5. FoldX energetic characterization of all models**

(A) Heatmap of FoldX alanine scan for all models. The color scale is confined to the limits [-1.6, 1.6]. Hotspot residues with a $\Delta\Delta G \geq 1.2$ or $\leq -1.2$ were marked. Positive and negative $\Delta\Delta G$ values indicate a destabilization and stabilization of the RAS effector models by alanine mutation at the indicated position, respectively.

(B) FoldX interaction energies average for the three representative structures on each target. The standard errors of the mean are displayed.

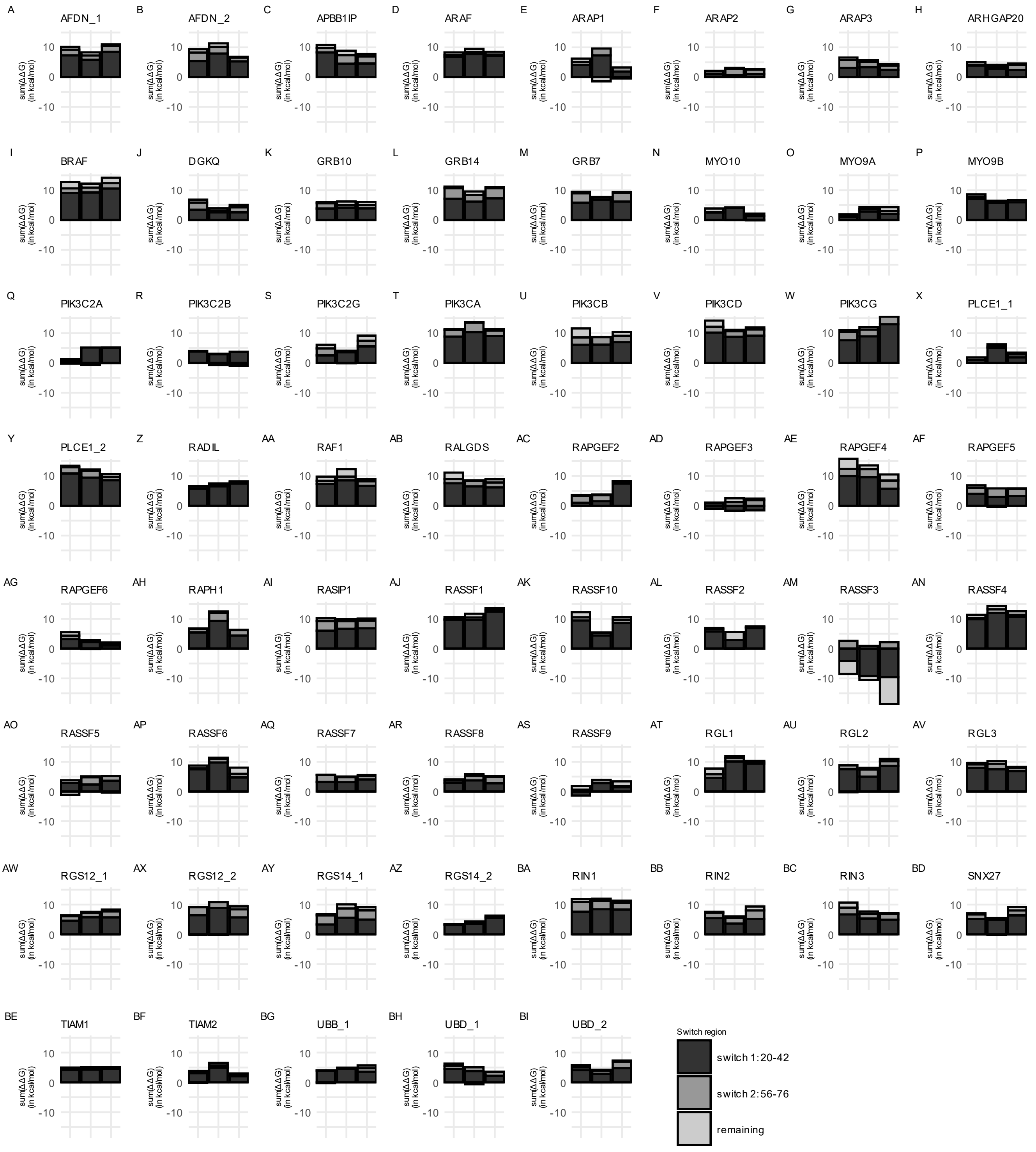


**Figure S6. Switch contributions of the three representative structures for each target**
For all modelled targets (A) – (BI), the individual switch contributions of the three representative models are visualized as bar charts.

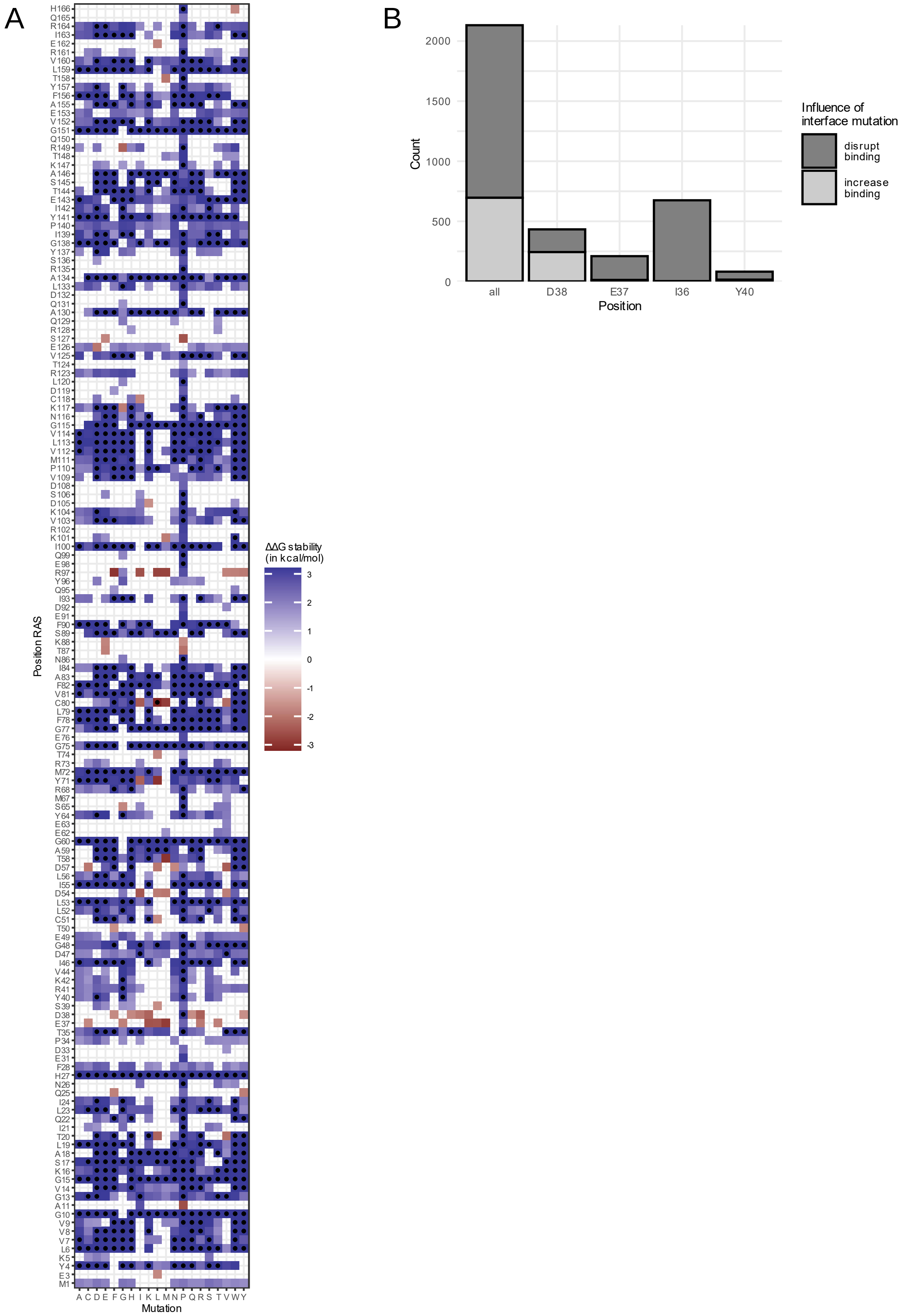


**Figure S7.**

A. Heatmap of FoldX stability of Ras proteins. The color scale is confined to the limits [-3.2, 3.2]. Hotspot residues with a ΔΔG ≥ 1.6 or ≤ -1.6 were marked. Positive ΔΔG values and negative ΔΔG values indicate a destabilization and stabilization of RAS, respectively.

B. Count of branch pruning mutations and their effect on the whole analysis and different hotspots.

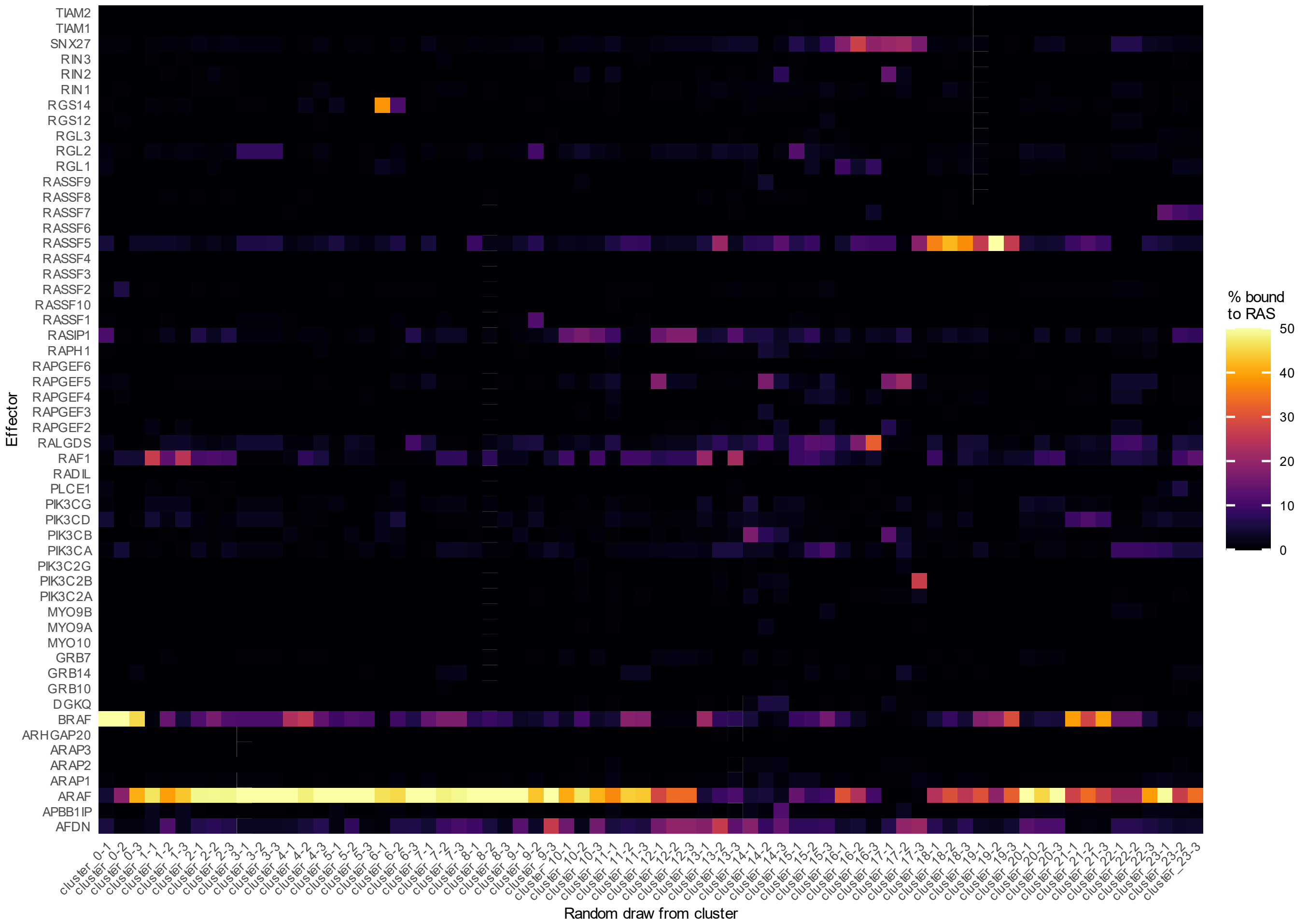


**Figure S8. Visualization of example system for the different UMAP-derived clusters.**
For each of the 19 clusters identified with OPTICS in the UMAP-transformed data, three members were randomly drawn, and the % of effectors bound to RAS was visualized. The colour scale restricted to [0 – 50] %.
