## Supplementary Tables for "Structure-based prediction of Ras-effector binding affinities and design of ‘branchegetic’ interface mutations"

**Supplementary Tables S1 to S5**

**Table S1.** Complex templates.

| Effector | Source | Identifier |
|---|---|---|
| PIK3CG | PDB | 1HE8 |
| RALGDS | PDB | 1LFD |
| PLCE1 | PDB | 2C5L |
| RASSF5 | PDB | 3DDC |
| RAF1 | PDB | 4G0N |
| GRB14 | PDB | 4K81 |
| AFDN | PDB | 6AMB |
| RGL1 | PDB | 7SCW |
| AFDN_1 | AlphaFold2 multimer | AF2c_ AFDN_1 |
| AFDN_2 | AlphaFold2 multimer | AF2c_ AFDN_2 |
| APBB1IP | AlphaFold2 multimer | AF2c_ APBB1IP |
| ARAF | AlphaFold2 multimer | AF2c_ ARAF |
| ARHGAP20 | AlphaFold2 multimer | AF2c_ ARHGAP20 |
| BRAF | AlphaFold2 multimer | AF2c_ BRAF |
| GRB10 | AlphaFold2 multimer | AF2c_ GRB10 |
| GRB14 | AlphaFold2 multimer | AF2c_ GRB14 |
| GRB7 | AlphaFold2 multimer | AF2c_ GRB7 |
| MYO9A | AlphaFold2 multimer | AF2c_ MYO9A |
| PIK3CA | AlphaFold2 multimer | AF2c_ PIK3CA |
| PIK3CB | AlphaFold2 multimer | AF2c_ PIK3CB |
| PIK3CG | AlphaFold2 multimer | AF2c_ PIK3CG |
| PIK3CD | AlphaFold2 multimer | AF2c_ PIK3CD |
| PLCE1_2 | AlphaFold2 multimer | AF2c_ PLCE1_2 |
| RADIL | AlphaFold2 multimer | AF2c_ RADIL |
| RAF1 | AlphaFold2 multimer | AF2c_ RAF1 |
| RALGDS | AlphaFold2 multimer | AF2c_ RALGDS |
| RAPGEF4 | AlphaFold2 multimer | AF2c_ RAPGEF4 |
| RAPGEF5 | AlphaFold2 multimer | AF2c_ RAPGEF5 |
| RAPH1 | AlphaFold2 multimer | AF2c_ RAPH1 |

| | | |
|---|---|---|
| RASIP1 | AlphaFold2 multimer | AF2c_ RASIP1 |
| RASSF1 | AlphaFold2 multimer | AF2c_ RASSF1 |
| RASSF2 | AlphaFold2 multimer | AF2c_ RASSF2 |
| RASSF3 | AlphaFold2 multimer | AF2c_ RASSF3 |
| RASSF4 | AlphaFold2 multimer | AF2c_ RASSF4 |
| RASSF5 | AlphaFold2 multimer | AF2c_ RASSF5 |
| RASSF6 | AlphaFold2 multimer | AF2c_ RASSF6 |
| RGL1 | AlphaFold2 multimer | AF2c_ RGL1 |
| RGL2 | AlphaFold2 multimer | AF2c_ RGL2 |
| RGL3 | AlphaFold2 multimer | AF2c_ RGL3 |
| RGS12_1 | AlphaFold2 multimer | AF2c_ RGS12_1 |
| RGS14_1 | AlphaFold2 multimer | AF2c_ RGS14_1 |
| RIN1 | AlphaFold2 multimer | AF2c_ RIN1 |
| RIN2 | AlphaFold2 multimer | AF2c_ RIN2 |
| RIN3 | AlphaFold2 multimer | AF2c_ RIN3 |

**Table S2.** Single templates. For each putative RBD, the AlphaFold2 prediction of the main UNIPROT sequence was downloaded from the AlphaFold Protein Structure Database (Release 1, accessed September 2021). The relevant domain was extracted either by using the domain information from PFAM, or for the proteins where PFAM had no RBD annotation, by using TMalign to identify the conserved ubiquitin superfold.

| **HGNC** | **UNIPROT ID** | **Residue start** | **Residue end** |
|---|---|---|---|
| ARAF | P10398 | 19 | 91 |
| BRAF | P15056 | 155 | 227 |
| RAF1 | P04049 | 56 | 131 |
| PIK3CA | P42336 | 187 | 289 |
| PIK3CB | P42338 | 194 | 285 |
| PIK3CG | P48736 | 217 | 309 |
| PIK3CD | O00329 | 187 | 278 |
| PIK3C2A | O00443 | 421 | 509 |
| PIK3C2B | O00750 | 375 | 463 |
| PIK3C2G | O75747 | 285 | 371 |
| RALGDS | Q12967 | 798 | 885 |
| RGL1 | Q9NZL6 | 648 | 735 |
| RGL2 | O15211 | 648 | 735 |
| RGL3 | Q3MIN7 | 613 | 700 |
| AFDN_1 | P55196_1 | 39 | 133 |
| AFDN_2 | P55196_2 | 246 | 348 |
| PLCE1_1 | Q9P212_1 | 2012 | 2114 |
| PLCE1_2 | Q9P212_2 | 2135 | 2238 |
| RIN1 | Q13671 | 624 | 706 |
| RIN2 | Q8WYP3 | 787 | 878 |
| RIN3 | Q8TB24 | 877 | 963 |
| SNX27 | Q96L92 | 273 | 362 |
| TIAM1 | Q13009 | 765 | 832 |
| TIAM2 | Q8IVF5 | 810 | 881 |
| ARHGAP20 | Q9P2F6 | 194 | 295 |
| ARAP1 | Q96P48 | 1172 | 1261 |
| ARAP2 | Q8WZ64 | 1326 | 1420 |

| | | | |
|---|---|---|---|
| ARAP3 | Q8WWN8 | 1117 | 1210 |
| DGKQ | P52824 | 395 | 494 |
| RASSF1 | Q9NS23 | 164 | 292 |
| RASSF2 | P50749 | 176 | 264 |
| RASSF3 | Q86WH2 | 79 | 186 |
| RASSF4 | Q9H2L5 | 174 | 262 |
| RASSF5 | Q8WWW0 | 236 | 364 |
| RASSF6 | Q6ZTQ3 | 218 | 306 |
| RASSF7 | Q02833 | 6 | 89 |
| RASSF8 | Q8NHQ8 | 1 | 82 |
| RASSF9 | O75901 | 25 | 119 |
| RASSF10 | A6NK89 | 4 | 133 |
| RAPGEF2 | Q9Y4G8 | 606 | 692 |
| RAPGEF3 | O95398 | 557 | 637 |
| RAPGEF4 | Q8WZA2 | 667 | 747 |
| RAPGEF5 | Q92565 | 240 | 321 |
| RAPGEF6 | Q8TEU7 | 749 | 835 |
| RASIP1 | Q5U651 | 144 | 259 |
| RADIL | Q96JH8 | 61 | 164 |
| APBB1IP | Q7Z5R6 | 176 | 263 |
| RAPH1 | Q70E73 | 269 | 355 |
| MYO9A | B2RTY4 | 14 | 112 |
| MYO9B | Q13459 | 15 | 114 |
| MYO10 | Q9HD67 | 1684 | 1791 |
| RGS12_1 | O14924_1 | 962 | 1032 |
| RGS12_2 | O14924_2 | 1034 | 1104 |
| RGS14_1 | O43566_1 | 302 | 373 |
| RGS14_2 | O43566_2 | 375 | 445 |
| GRB7 | Q14451 | 100 | 186 |
| GRB10 | Q13322 | 166 | 250 |
| GRB14 | Q14449 | 106 | 192 |
| UBB_1 | P0CG47_1 | 1 | 76 |

| | | | |
|---|---|---|---|
| UBD_1 | O15205_1 | 6 | 81 |
| UBD_2 | O15205_2 | 90 | 163 |

**Table S3.** The hyperparameter space explored for the selection of representative structures. The best combination of hyperparameters is marked in **bold text**. For feature selection, hotspots were ranked by number of occurrences in all models for a specific target. min_samples, min_cluster_size and xi are hyperparameters of the OPTICS algorithm. For more information, please refer to the documentation: https://scikit-learn.org/stable/modules/generated/sklearn.cluster.OPTICS.html

| Hyperparameter | Options |
|---|---|
| Feature selection strategy | all, only hotspots, **top 10 hotspots + interaction**, top 5 hotspot + interaction, top 3 hotspots + interaction |
| Dimensionality reduction | **None**, PCA, ICA, UMAP, kPCA_rbfdot, kPCA_polydot, kPCA_anovadot |
| Number of components after dimensionality reduction | 3, 5, 7 |
| min_samples | 3, 4, **5**, 7, 10 |
| min_cluster_size | 5, **15**, 30, 50, 70 |
| xi | 0.01, 0.015, 0.02, 0.025, 0.03, 0.035, 0.04, 0.045, 0.05, 0.055, 0.06, 0.065, **0.07** |
| Selection of representative structures | Lowest energy cluster top 3 structures by FoldX interaction energy, **lowest energy cluster top 3 structures by smallest average distance** |

**Table S4.** The hyperparameter space explored for the regression of binding energies based on FoldX-derived features. The best combination of hyperparameters is marked in **bold text**. For all regressors, a two-step pipeline was used. In step 1, feature selection was performed using SelectKBest. In the step 2, the hyperparameter space for different regressors was explored. For more information, please refer to the documentation: https://scikit-learn.org/stable

| Step | Model | Hyperparameter | Options |
|---|---|---|---|
| 1 | SelectKBest | k | 1:27; best is **10** |
| | | score_func | **mutual_information_regression,** f_regression |
| 2 | Lasso | alpha | 0:1 in 1000 steps |
| | LassoLars | alpha | 0:1 in 1000 steps |
| | Ridge | alpha | 0:1 in 1000 steps |
| | ElasticNet | alpha | 0:1 in 1000 steps |
| | | l1_ratio | 0.005, 0.01, 0.1, 0.5, 0.7, 0.9, 0.95, 0.99, 1 |
| | LinearRegression | - | - |
| | SVR | kernel | linear |
| | | C | 0.1, 1, 10, 100, 1000, 10000 |
| | | epsilon | 0.001, 0.01, 0.1, 0.5, 1 |
| | **SVR** | kernel | poly, **rbf**, sigmoid |
| | | C | 0.1, 1, 10, 100, **1000**, 10000 |
| | | epsilon | 0.001, 0.01, 0.1, **0.5**, 1 |
| | | gamma | 1 0.1, 0.01, **0.001**, 0.0001 |
| | RandomForestRegression | n_estimators | 50, 75, 100, 125 |
| | | max_depth | None, 2, 3, 4, 5, 6 |
| | | max_features | 0.1, 0.2, 0.3, 0.4, 0.5, 0.6, 0.8, 1 |
| | | bootstrap | True, False |

**Table S5.** Predicted $K_d$ values in comparison with experimental $K_d$ values. nb stands for not binding.

| **Target** | **Predicted $K_d$ (uM)** | **Experimental $K_d$ (uM)** | **PMID** | **Used in test or training set** |
|---|---|---|---|---|
| ARAF | 0.257664 | 0.07 | 24441586 | Training |
| BRAF | 0.020433 | 0.04 | 24441586 | Training |
| RAF1 | 0.233102 | 0.08 | 11292826 | Test |
| PIK3CA | 0.828134 | | | |
| PIK3CB | 5.106858 | | | |
| PIK3CG | 1.012316 | 2.9 | 11136978 | Training |
| PIK3CD | 0.986412 | | | |
| PIK3C2A | 122.4118 | | | |
| PIK3C2B | 588.1618 | | | |
| PIK3C2G | 17.89329 | | | |
| RALGDS | 2.95131 | 1 | 11292826 | Training |
| RGL1 | 6.076352 | 3.5 | 9753431 | Test |
| RGL2 | 3.744253 | 0.09 | 9753431 | Training |
| RGL3 | 5.635062 | | | |
| AFDN_1 | 3.096808 | 3.03 | 11292826 | Training |
| AFDN_2 | 2.633623 | | | |
| PLCE1_1 | 36.95527 | | | |
| PLCE1_2 | 1.137651 | 0.82 | 15826668 | Test |
| RIN1 | 1.577532 | 0.88 | 15826668 | Training |
| RIN2 | 11.32005 | 11 | 15826668 | Test |
| RIN3 | 19.36871 | | | |
| SNX27 | 25.84413 | | | |
| TIAM1 | 45.29575 | | | |
| TIAM2 | 107.8572 | | | |
| ARHGAP20 | 45.16912 | | | |
| ARAP1 | 15.27132 | | | |
| ARAP2 | 42.52022 | | | |
| ARAP3 | 23.03746 | | | |

| | | | | |
|---|---|---|---|---|
| DGKQ | 18.45376 | | | |
| RASSF1 | 13.29281 | 39 | 15826668 | Training |
| RASSF2 | 66.08577 | 147 | 33930461 | Training |
| RASSF3 | | 500 | 33930461 | |
| RASSF4 | 212.0512 | 193 | 33930461 | Training |
| RASSF5 | 1.451177 | 0.21 | 24441586 | Training |
| RASSF6 | 72.1179 | 91 | 33930461 | Training |
| RASSF7 | 15.30952 | 140 | 33930461 | Training |
| RASSF8 | 17.0793 | nb | 33930461 | |
| RASSF9 | 135.0758 | 179 | 33930461 | Training |
| RASSF10 | 32.53544 | nb | 33930461 | |
| RAPGEF2 | 14.19324 | 33 | 15826668 | Training |
| RAPGEF3 | 76.75141 | nb | 15826669 | |
| RAPGEF4 | 2.465967 | | | |
| RAPGEF5 | 9.242346 | | | |
| RAPGEF6 | 103.6208 | | | |
| RASIP1 | 2.652572 | nb | 15826668 | |
| RADIL | 49.68458 | | | |
| APBB1IP | 17.66283 | | | |
| RAPH1 | 17.34223 | | | |
| MYO9A | 41.23323 | | | |
| MYO9B | 88.87451 | nb | 15826669 | |
| MYO10 | 71.10372 | | | |
| RGS12_1 | 9.043251 | | | |
| RGS12_2 | 7.166583 | | | |
| RGS14_1 | 7.415029 | 14 | 24441586 | Training |
| RGS14_2 | 30.27894 | | | |
| GRB7 | 4.097531 | | | |
| GRB10 | 51.39415 | | | |
| GRB14 | 4.328095 | 3.6 | 26423700 | Training |
| UBB_1 | 33.13215 | nb | 16310215 | |
| UBD_1 | 52.91385 | nb | 16310215 | |

| | | | |
|---|---|---|---|
| UBD_2 | 18.4631 | nb | 16310215 |
